# Regulation of pitcher fluid volume and properties in six ecologically distinct Bornean *Nepenthes* species

**DOI:** 10.64898/2026.06.10.731308

**Authors:** C.N.S. Andrew, J-Y. Bu, I.J. Howell, F. Metali, T.U. Grafe, W. Federle

**Affiliations:** Department of Zoology, University of Cambridge, UK, CB2 3EJ; Department of Mechanical Engineering, National University of Singapore, Singapore, 117575; Faculty of Science, Universiti Brunei Darussalam, Brunei

## Abstract

Carnivorous *Nepenthes* plants capture and digest prey in fluid-filled pitchers, whose fluid level is subject to fluctuations caused by rain and evaporation. It was found that *N. rafflesiana* pitchers can reduce these fluctuations by regulating their fluid volume and composition. Here, we investigate whether and to what extent this ability occurs in other *Nepenthes* species varying in morphology, trapping mechanisms, habitat, lifespan and nutrient sources. We hypothesised that species more exposed to fluid level fluctuations or relying more on fluid for prey capture would exhibit more intensive regulation. In six Bornean *Nepenthes* species, we quantified the effect of pitcher fluid level on prey-capture efficiency, the natural fluctuations in fluid volume, and the plants’ response to manipulations of the fluid’s volume, concentration and pH.

The prey capture of *Nepenthes* species depended to varying degrees on the pitcher fluid level: in species with waxy inner walls, capture efficiency decreased at high fluid levels, as prey could escape easily. In contrast, species with wax-free walls and steep peristomes retained prey efficiently, regardless of fluid level. Although all *Nepenthes* species were able to secrete and absorb water and electrolytes, the regulation rate varied considerably. The rates of the various secretion and absorption processes (concentration/volume-dependent rates of electrolyte/water transport) were positively correlated with one another, suggesting that they all depend on the activity of the pitcher glandular epithelium. While differences and similarities in fluid regulation may be partly due to phylogeny, the observed species differences can also be explained by the pitchers’ trapping mechanisms and ecological factors. We found weaker regulation in *Nepenthes* species growing in sheltered understorey habitats, and in species utilising alternative nutrient sources, which may reduce their reliance on pitcher fluid. Our study highlights the diversity of ecological strategies in pitcher plants and demonstrates how these are influenced by environmental conditions.

**Lay Summary:** *Nepenthes* pitcher plants have fluid-filled pitchers to capture and digest prey. Can they control the volume and composition of their fluids? We found that all six species studied regulated their fluids, but to varying degrees. Species differences can be explained by the plants’ habitat, pitcher morphology and nutrient acquisition strategies.

## 1. Introduction

As climate change leads to increasingly extreme weather conditions worldwide, many ecosystems are in immediate jeopardy. At the heart of the climate crisis lies the unpredictability of global water availability. Water stress imperils plants and the ecosystems they support (Bartholomeus *et al*., 2011), as plants require water for photosynthesis, nutrient transport, turgor pressure and transpiration. Beyond these general functions of water, openly accessible watery fluids are important for many plant-animal interactions. Bromeliads, for example, support whole ecosystems in fluid-filled hollows between their leaves (Frank, 1983; Gerlach, 2011). Weather fluctuations threaten to flood or evaporate such plant-held fluid reservoirs, putting these plant-animal interactions at risk (Kitching, 2000; Andrew *et al*., 2026). How are these exposed fluid pools affected by weather conditions and do plants regulate these fluids in order to maintain their function?

Plants that use open fluids include a range of taxonomically disparate carnivorous species. Their fluids are essential for capturing and digesting prey and include sticky droplets suspended on the hairs of *Drosera*, *Roridula*, *Pinguicula* and *Byblis*, digestive fluids secreted by Venus flytraps or the fluids contained in the pitchers of *Nepenthes*, *Sarracenia*, *Heliamphora* or *Cephalotus*.

*Nepenthes* plants possess leaves modified into pitchers that serve as pitfall traps to capture arthropod prey. Insect visitors are attracted using odour and visual signals (Di Giusto *et al*., 2008; Bennett and Ellison, 2009) and trapped when they fall from the slippery pitcher rim (peristome) (Bohn and Federle, 2004) or the slippery underside of the springboard-like lid in some species (Bauer *et al*., 2012b; Lenz and Bauer, 2022). The prey falls into the fluid that partially fills the pitcher (Clarke, 1997) and is prevented from escape in many species by a slippery waxy inner pitcher wall (Gaume *et al*., 2004). The over 200 *Nepenthes* species identified to date (McPherson, 2023) show great variation in terms of their habitat, prey composition, and associations with animals, as well as their pitcher morphology, pitcher fluid properties, and capture mechanisms (Bauer *et al*., 2012a; Moran *et al*., 2013).

The function of the fluid contained within *Nepenthes* pitchers is to immobilise and digest captured prey. Its composition and volume are likely affected by rain and evaporation. Pitcher fluids vary from watery to sticky across species (Bonhomme *et al*., 2011; Kang *et al*., 2021) and differ in their efficiency in retaining live prey in the pitcher (Bauer *et al*., 2011). The pitcher fluid of *N. rafflesiana* has a lower surface tension than water, and due to its viscoelastic nature, is also resistant to dewetting, so prey becomes more easily coated and submerged in the pitcher fluid (Kang *et al*., 2021). We recently showed that prey capture decreases in *N. rafflesiana* when pitcher fluid levels are very high after rainfall, or low following evaporation (Andrew *et al*., 2026). Prey retention is also reduced when the pitcher fluid is strongly diluted (Gaume *et al*., 2007). Since changes in fluid volume and composition can be detrimental to prey capture, how can pitchers maintain their open fluid pools under fluctuating weather conditions? *N. rafflesiana* has a remarkable ability to maintain an intermediate fluid level in its pitchers by absorbing or secreting fluid, thereby maximising prey capture and nutrient gain (Andrew *et al*., 2026).

*Nepenthes* species may be more or less exposed to fluid level fluctuations under natural conditions due to their varying pitcher morphology or habitat conditions, or they may be less reliant on their pitcher fluids for prey capture due to specialised nutrient acquisition strategies. Do all *Nepenthes* species regulate pitcher fluid volume in the same way as *N. rafflesiana*? *Nepenthes* species unable to respond to weather-related changes to their fluid may be at greater risk under extreme weather patterns resulting from climate change. Climate change is already projected to threaten *Nepenthes* species due to habitat loss (Gray *et al*., 2017) and it could impact humidity-dependent capture mechanisms in some species (Bohn and Federle, 2004; Bauer *et al*., 2008; Moran *et al*., 2013; Bauer *et al*., 2015). However, the effects of weather fluctuations on the efficiency of *Nepenthes* prey capture mechanisms have yet to be investigated.

### Factors relevant to pitcher fluid regulation

Various factors may influence how beneficial it is for pitcher plants to regulate their fluid. For example, plant habitat and pitcher morphology can influence the exposure of pitchers to fluid level fluctuations. Other pitcher traits, as well as ecological factors including alternative strategies for nutrient uptake, the pitchers’ lifespan, and infauna and digestion may influence the extent to which pitcher plants actually require their fluid.

#### Habitat

*Nepenthes* pitcher plants grow on nutrient-poor soils in kerangas, peat-swamp and montane forests. These habitats differ in their exposure to rain and dry conditions, and thus the risk of pitcher flooding and fluid evaporation. Protection by tree canopies and foliage may prevent flooding and excessive evaporation (Pickering *et al*., 2021). It has been shown that *Nepenthes* species with wetness-dependent peristome and fluid-based capture mechanisms are much more restricted to humid environments than species that trap prey with slippery waxy surfaces (Moran *et al*., 2013).

#### Pitcher morphology

*Nepenthes* pitchers differ in the size of the pitcher lids relative to the pitcher mouth and in the lids’ angles of inclination (Chomicki *et al*., 2024), both affecting the influx of rain and consequently fluid level fluctuation. Pitcher morphology also mediates different prey capture mechanisms which depend to varying degrees on the pitcher fluid. *Nepenthes* capture mechanisms can be categorised as peristome-trapping, wax-trapping and fluid-retention (Moran *et al*., 2013), with many species utilising more than one of these strategies. Pitcher traits such as a wider and steeper peristome (Bauer *et al*., 2012a; Moulton *et al*., 2023) or a slippery wax crystal coating (Gaume *et al*., 2004; Gorb *et al*., 2005; Riedel *et al*., 2007), ensure efficient prey retention so that prey retention via the fluid may be less important. Pitcher fluid regulation may be more important in species whose trapping mechanism relies primarily on fluid retention.

#### Specialised nutrient acquisition strategies

While most *Nepenthes* species primarily capture live insect prey, some species have evolved other nutrient acquisition strategies, which may reduce the need for the pitcher fluid to contribute to prey capture. For example, the funnel shape and widened mouth of several *Nepenthes* species is a specialisation to capture shrew excrement (Chin *et al*., 2009; Clarke *et al*., 2009); the wide mouth and reduced lid of *N. ampullaria* facilitate the capture of falling detritus (Cresswell, 1998; Moran *et al*., 2003; Pavlovic *et al*., 2011; Gilbert *et al*., 2022). Some *Nepenthes* have specific associations with animals which are essential for the plants’ nutrient gain (Moran *et al*., 2001; Merbach *et al*., 2002; Grafe *et al*., 2011). For example, the narrow pitchers with a sudden taper of *N. hemsleyana* (previously known as *N. rafflesiana* var. *elongata* or *N. baramensis* (Scharmann and Grafe, 2013)) attract roosting Hardwicke’s Woolly bats that provide pitchers with nutrients via their excrement (Grafe *et al*., 2011; Schöner *et al*., 2013; Lim *et al*., 2014; Schöner *et al*., 2015a; Schöner *et al*., 2015b) and *Polyrhachis nepenthicola* ants use *N. stenophylla* pitchers as a nesting spot (Grafe and Kohout, 2013). These associations provide pitchers with nutrients via waste products so that the pitcher fluid may be less essential. Other *Nepenthes* species are host to predators that ambush prey, potentially increasing the pitcher’s nutrient gain and reducing the need for the pitcher fluid to contribute to prey capture, such as *Misumenops nepenthicola* and *Thomisus nepenthiphilus* crab spiders found in *N. gracilis* and *N. rafflesiana* pitchers (Lam *et al*., 2019b; Karl and Bauer, 2020). In all these species, the pitcher fluid is not (or to a lesser extent) needed to retain moving insect prey, potentially reducing the need for fluid regulation.

#### Pitcher lifespan

Pitcher lifespan varies strongly among the six *Nepenthes* species studied, ranging from ∼2 months to over 2.5 years (Osunkoya *et al*., 2008; Thornham *et al*., 2012). Fluid volume fluctuations occur on much shorter timescales, with rainfall in a single day significantly altering the pitcher fluid level (Andrew *et al*., 2026). It is possible that shorter-lived pitchers prioritise efficient prey capture in a short timespan in order to lower the payback time per pitcher, and thus invest more into fluid regulation mechanisms. In contrast, longer-lived pitchers can capture prey over a longer period of time and therefore may not need to invest as much into capture efficiency. However, the construction costs for longer-lived pitchers are higher (Osunkoya *et al*., 2008), which increases the required payback. It is therefore possible that pitchers trade off pitcher construction costs with costs for more efficient prey capture (including via fluid regulation).

#### Infauna and digestion

Most *Nepenthes* pitchers host complex communities of microorganisms and arthropods which play an important role in the digestion of prey (Clarke and Kitching, 1993; Adlassnig *et al*., 2011; Lam *et al*., 2017; Ellison and Adamec, 2018; Gaume *et al*., 2019; Lam *et al*., 2019b; Lam *et al*., 2020). These diverse communities enhance the availability of nutrients to the plant (Lam *et al*., 2017; Leong *et al*., 2018; Lam *et al*., 2019a). A sufficient volume of pitcher fluid is required to sustain these infaunal communities and for the pitchers’ digestion process as a whole. Thus, regulation to maintain the volume of pitcher fluid is likely beneficial for all *Nepenthes* species in order to maintain efficient digestion.

#### Hypothesis and aims

We hypothesise that *Nepenthes* species that are more exposed to fluid fluctuations and more dependent on their pitcher fluid for prey capture and digestion show more intensive fluid regulation. To test this hypothesis, we investigated in six *Nepenthes* species native to Brunei, Borneo (Fig. 1 and Table 1) how much pitcher fluid volumes fluctuate under natural conditions, how pitcher fluid level influences prey capture efficiency and to what extent the pitchers are able to regulate the fluid volume and composition.

**Figure 1:**
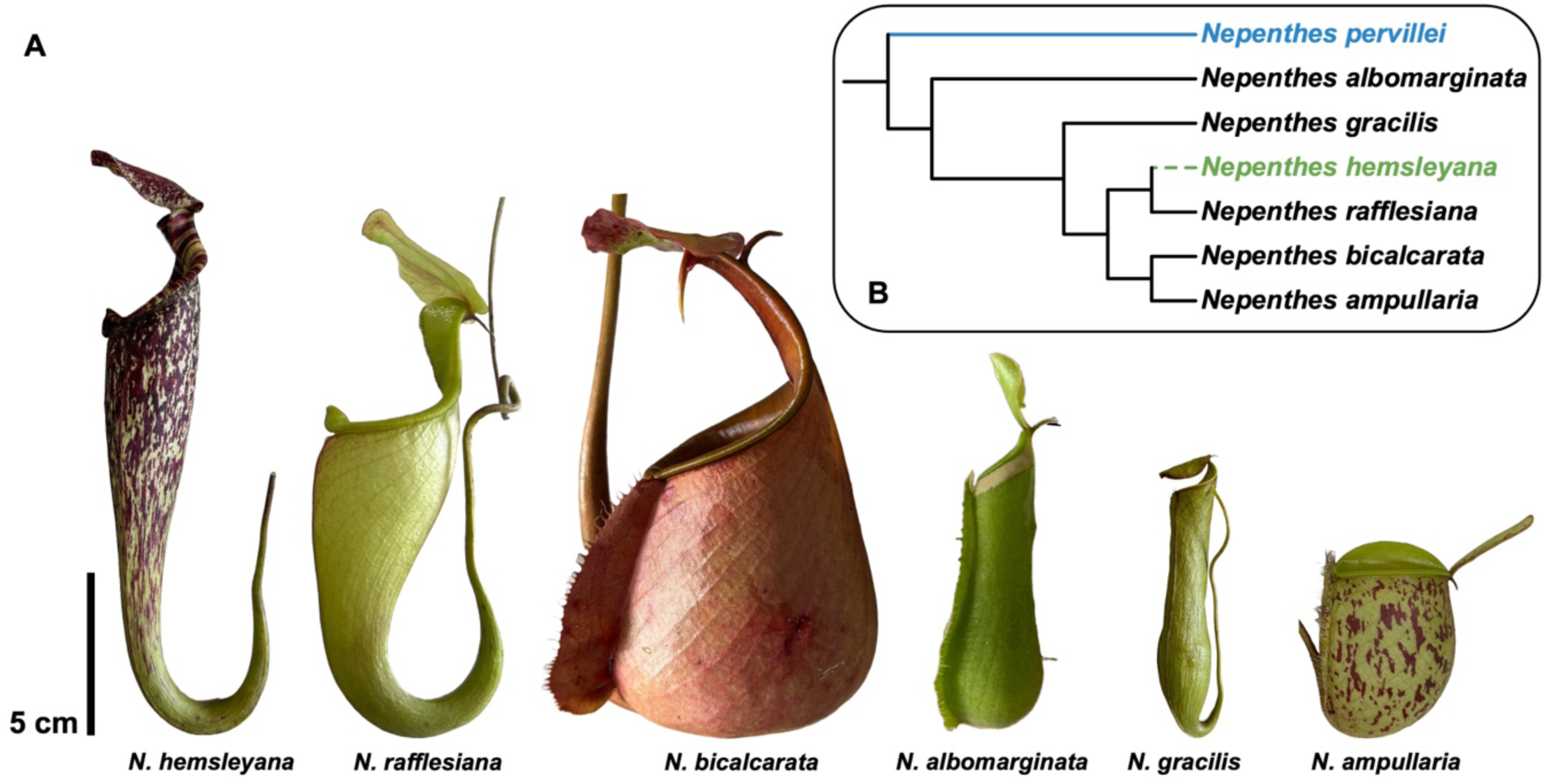
(A) Morphology of the six Bornean *Nepenthes* species used in this study. All pitchers were collected from their respective field sites and photographed within a few hours in the laboratory. (B) Phylogeny of the species used in this study, based on tRNA sequences retrieved from NCBI (Meimberg *et al*., 2006) with *Nepenthes pervillei* (blue) included as an outgroup. *Nepenthes hemsleyana* (green) was grafted onto the tree as a sister species of *Nepenthes rafflesiana* (Scharmann and Grafe, 2013; Murphy *et al*., 2020). Branch lengths are not scaled to denote genetic divergence times as no sequences were available for *N. hemsleyana*. **Alt text:** (A) Photographs of the side profile of six *Nepenthes* pitchers of different species. (B) A phylogenetic tree depicting the relatedness of the six *Nepenthes* species pictured.

**Table 1:**
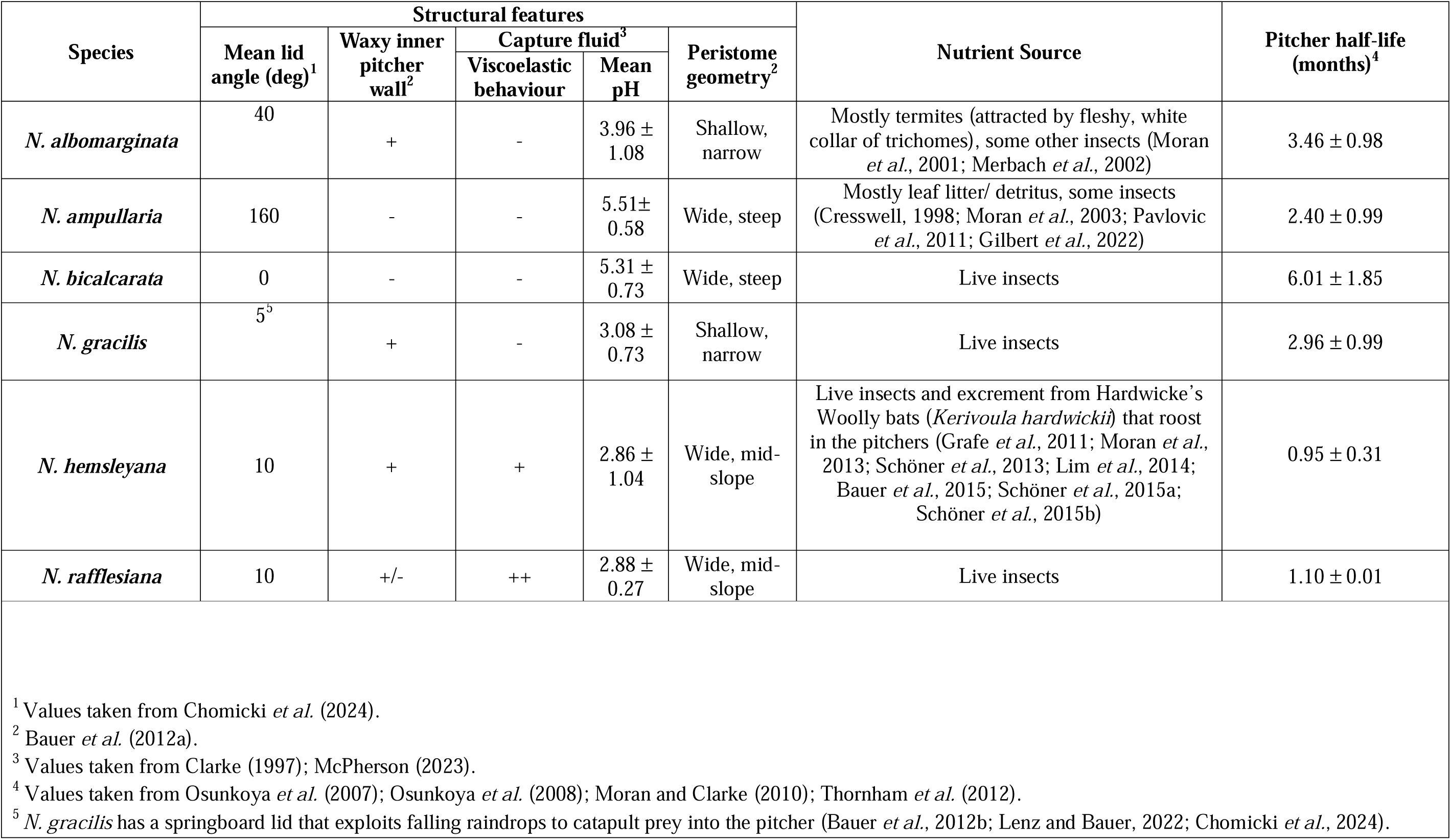
Comparison of the main features of the six Bornean *Nepenthes* species used in this study. Mean lid angle, wax inner lining and peristome data are obtained from literature while capture fluid parameters are based on my own experiments. In combination, these features describe the main prey capture and nutrient acquisition strategies of the species listed and may point to some species being more reliant on the properties of their capture fluid than others.

## 2. Materials and Methods

All experiments were performed on *Nepenthes* plants in their natural habitats in Brunei (Borneo) from June-October 2024. *Nepenthes rafflesiana*, *Nepenthes hemsleyana* and *Nepenthes gracilis* were studied at a site of degraded kerangas (heath) forest (4°44’23.9“N 114°35’36.4”E), *Nepenthes ampullaria* and *Nepenthes albomarginata* in secondary heath forest (4°35’21.0“N 114°30’16.1”E), and *Nepenthes bicalcarata* in peat swamp forest (4°33’34.9“N 114°29’19.6”E). Fieldwork was carried out under forest entry permit no. JPH/UND/17.

*Nepenthes* plants generally produce two morphologically distinct trap types: ‘upper’ pitchers grow above ground on mature vines and possess a funnel-like morphology while ‘lower’ pitchers are found at ground level and are more bulbous in shape. Both pitcher types attract, capture and digest prey, and they can differ in their prey composition but their fundamental function remains the same (Juniper *et al*., 1989; Moran, 1996; Clarke, 1997). For this reason, the pitcher type used for each species was selected based on abundance and availability (upper pitchers are rare for some species), namely upper pitchers for *N. rafflesiana*, *N. hemsleyana* and *N. gracilis* and lower pitchers for *N. ampullaria*, *N. albomarginata* and *N. bicalcarata*.

For each of the six species studied, 60 pitchers were investigated, each from different plants. As insect capture rates are higher in more recently opened pitchers (Bauer *et al*., 2009), we ensured that all pitchers were ‘fresh’ by selecting only pitchers from the most recently developed internodes that had a good overall appearance, with a bright green peristome colour and clear pitcher fluid.

In all time series experiments, the observation period lasted for 12 days with daily measurements taken on days 0-5 and a final measurement on day 12.

### 2.1 Measurements of naturally occurring pitcher fluid properties

The physical and chemical properties of the pitcher fluids of each species were studied by taking ten 2-ml fluid samples per species. The chemical properties, conductivity, pH and K^+^ concentration were measured *in situ*. Each of the fluid properties were measured using portable meters (conductivity: LAQUAtwin conductivity meter EC-33, pH: LAQUAtwin pH meter PH-33, K+: LAQUAtwin potassium ion meter K-11, HORIBA Ltd., Kyoto, Japan). Between field measurements, the meters were rinsed with distilled water until their displays showed <10 μS/cm, pH 7 (± 0.1) and <10 ppm, respectively.

The fluid’s surface tension and extensional relaxation time were measured in the laboratory at 25°C within 3 hours of collection for 5 samples per species. Surface tension was measured using a portable pendant drop tensiometer fabricated from 3D printed parts. Approximately 0.5ml of fluid was drawn into a 2-ml syringe fitted with a 1.8 mm diameter needle with a flattened tip. Fluid droplets were produced at the needle tip by pumping using a screw mechanism. The droplets were recorded with a high-speed camera (Ximea MQ003MG-CM, Ximea GmbH, Slovakia) and images were analysed using OpenDrop (Huang *et al*., 2021); we used 1.2 kg/m³ as the density of air and 1000 kg/m³ as the density of pitcher fluid (identical to the density of water, determined using a portable density meter on 7 pitcher fluid samples, 1000 ± 1 kg/m^3^).

To measure the extensional relaxation time of the fluid, a measure of its viscoelastic behaviour, we used a portable extensional rheometer (Collett *et al*., 2015) which rapidly stretches a small fluid sample between two metal pistons. Using a micropipette, 3μl of pitcher fluid was placed between the rheometer’s pistons (1.2mm diameter) before a battery-operated solenoid rapidly moved one piston away, stretching the fluid into a filament. The fluid filament was filmed with a high-speed camera (Ximea MQ003MG-CM, Ximea GmbH, Slovakia) at 300fps. The decay in fluid filament diameter over time was analysed using a custom-built MATLAB GUI to fit an exponential curve to the data. Relaxation time was then calculated using the Upper Convected Maxwell model as in Collett *et al*. (2015).

### 2.2 Measurement of pitcher fluid level

To investigate the naturally occurring fluid levels in the pitchers of each species, 10 pitchers per species were selected and fluid levels were measured on all sampling days. Fluid levels were measured by shining a torch on the pitcher from behind and marking the fluid level with a pen to measure its height (L) above the tendril base using a calliper (Andrew *et al*., 2026). The pitcher height (H) (tendril base to peristome) was recorded to allow the calculation of relative fluid level (L/H). At the end of the pitcher fluid level measurements, fluid volumes were determined from the recorded fluid levels by pipetting water in 0.1 mL increments up to each level mark on the pitcher.

The rainfall during the experimental period was recorded using a rain gauge (Rain Collector II 7395-024, Davis Instruments, California, USA; resolution: 0.22 mm/count) connected to a datalogger (TGP-4901, Gemini Data Loggers, Chichester, UK) that measured cumulative precipitation at each of the field sites in intervals of 5 minutes.

### 2.3 Effect of pitcher fluid level on prey capture efficiency

A single representative pitcher along with enough fresh pitcher fluid to fill the pitcher was collected for each species and ∼500 *Dolichoderus cuspidatus* ants from the same colony were collected and housed in a Fluon-coated container. To study the effect of pitcher fluid level on prey capture efficiency, the pitchers were successively filled with pitcher fluid up to 25, 50, 75 and 100% levels and the peristome was wetted with distilled water. The pitchers were placed one at a time into the box containing the ants which were allowed to climb freely over the pitcher. For the first 20 ants that slipped and fell from the peristome, the outcome (captured or escaped within 5 minutes) was recorded, and these ants were then removed for subsequent trials.

### 2.4 Response to changes in pitcher fluid volume

To investigate whether pitchers from different species show a response to changes in their fluid level, we selected two sets of 10 pitchers per species from different plants and either (a) removed all their fluid or (b) filled them to the peristome with fluid from other pitchers of the same species and age. Pitcher height, fluid level, conductivity and pH were measured before experimental manipulation and in the case of fluid addition, the same measurements were taken again directly after treatment. All pitchers were then covered with a polypropylene sheet folded into a cone attached to the pitcher’s tendril to exclude any influx of rainwater during the experimental period. On all sampling days the fluid level and fluid properties were recorded again for each pitcher.

### 2.5 Response to changes in pitcher fluid concentration

We studied how pitchers respond to changes in the ionic concentration of the pitcher fluid. We selected two sets of 10 pitchers per species and measured their initial fluid level and fluid properties before draining, rinsing and refilling them with either (a) distilled water or (b) 100 mM KCl up to their original fluid level. Pitchers were covered as before, and fluid level and properties (conductivity and pH) were recorded immediately after experimental manipulation and on each subsequent sampling day.

### 2.6 Response to changes in pitcher fluid pH

We tested whether the pitchers of each species can respond to the alkalisation of their pitcher fluid via the secretion of protons. Using a micropipette, 0.1M NaOH were added in 20μL increments and the fluid was thoroughly mixed until a pH of 11 ± 0.5 was reached. Pitchers were covered as before, and fluid level and properties were recorded both before the treatment and on each sampling day.

## 3. Results

### 3.1 Pitcher fluid level

The average pitcher fluid levels of the six *Nepenthes* species ranged from 33.4 to 69.5% (Fig. 2). The species’ fluid levels differed significantly (ANOVA: F_5,54_=26.0, p<0.001), with the lowest fluid levels in *N. hemsleyana* and *N. gracilis*, and highest in *N. bicalcarata* and *N. rafflesiana* (Fig. 2, Table S1).

**Figure 2:**
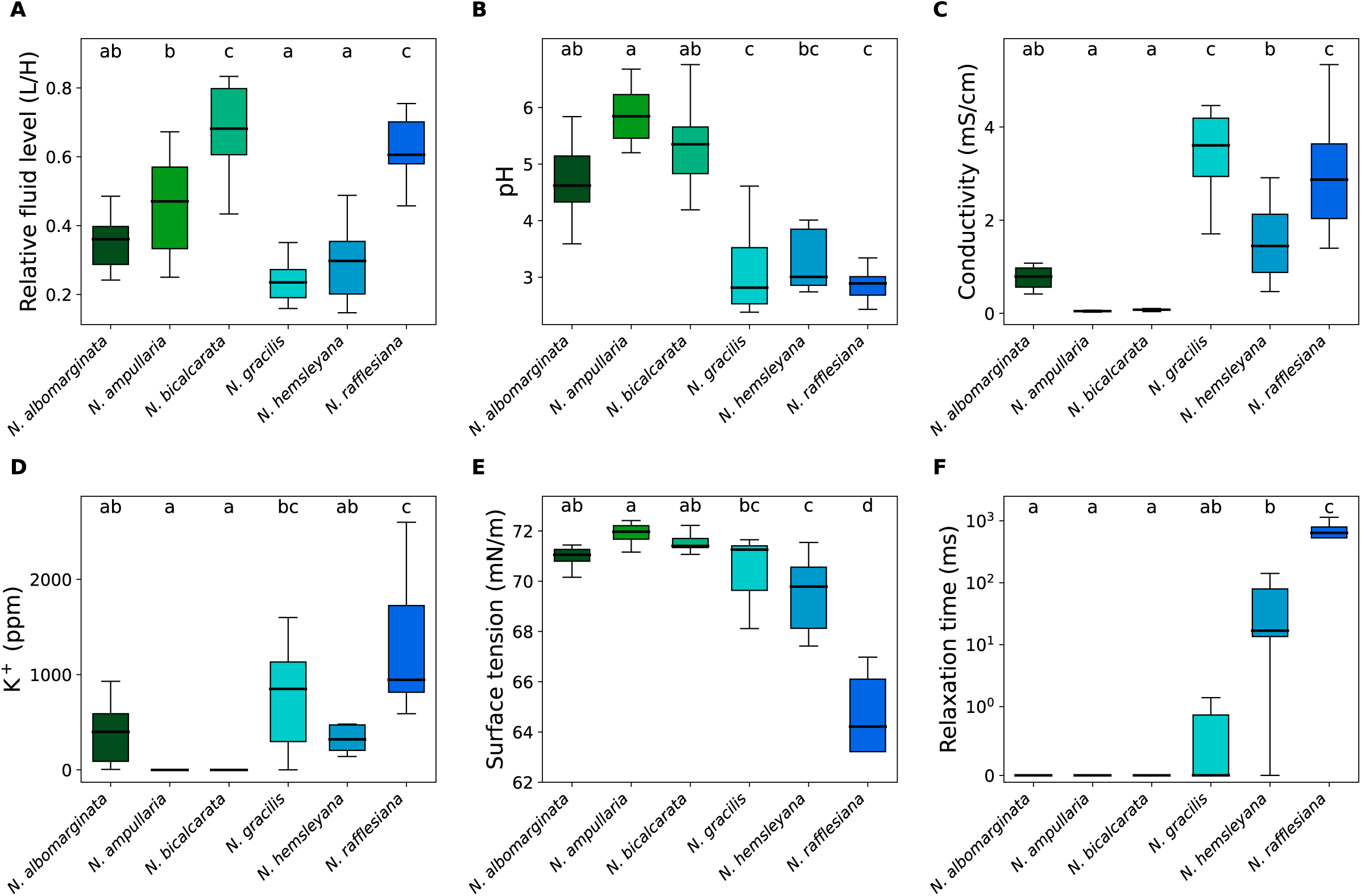
Physical and chemical properties of pitcher fluids in six *Nepenthes* species. For each species, 10 fresh pitchers from separate plants were measured in the field. (A) Relative fluid level (fluid level L/pitcher height H) (colour-filled boxes), (B) pH, (C) Conductivity, (D) Potassium (K^+^) concentration, (E) Surface tension and (F) Extensional relaxation time of the pitcher fluids found in each of the six species. The plot shows medians (centre lines), interquartile ranges (boxes) and the largest and smallest values (whiskers) that are not outliers. Letters a-d represent the statistical significance of differences between species, based on ANOVA (A & B), Welch-ANOVA (C, D & E) or Kruskal-Wallis (F) (all p<0.01) and Tukey’s HSD (A & B), Games-Howell (C, D & E) or Dunn’s (F) post-hoc tests. **Alt text:** Boxplots comparing pitcher fluid traits of each of the six *Nepenthes* species with letters denoting the result of pairwise comparisons.

Pitcher fluid levels showed statistically significant daily fluctuations (Table 2). On day 1, we observed a significant fluid level increase in *N. gracilis* despite no rainfall, indicating fluid secretion by the pitchers (Table 2). Significant fluid level decreases during dry days (likely due to evaporation) were observed in *N. albomarginata* and *N. bicalcarata* (Table 2). A significant fluid level decrease on a day with some rainfall (suggesting fluid absorption) was observed in *N. hemsleyana* (Table 2).

**Table 2:** One-sample t-tests comparing fluid levels of 10 pitchers per species between consecutive days, indicating significant fluid level changes (Fig. 3). All significant (non-zero) fluid level changes are shown. Rows coloured pink indicate days where there was a significant reduction in fluid level, indicative of pitcher fluid evaporation or absorption. Green rows indicate an increase in fluid level despite no rainfall, evidence for fluid secretion. Blue rows are significant fluid level increases during rainfall, indicative of rainwater influx.

| Species | Day | Rainfall /mm | Mean Fluid Level Change /mm | t<sub>9</sub> | p |
| --- | --- | --- | --- | --- | --- |
| *N. albomarginata* | 2-3 | 0.0 | -2.5 | -2.2 | 0.048 |
| *N. bicalcarata* | 0-1 | 0.0 | -4.1 | -2.3 | 0.044 |
| *N. gracilis* | 0-1 | 0.0 | 4.8 | 3.0 | 0.015 |
| *N. hemsleyana* | 0-1 | 10.12 | 6.5 | 5.8 | 0.000 |
|  | 1-2 | 19.58 | 6.5 | 2.3 | 0.044 |
|  | 2-3 | 6.60 | -7.7 | -3.4 | 0.007 |
| *N. rafflesiana* | 0-1 | 11.88 | 2.5 | -2.2 | 0.048 |
|  | 4-5 | 0.0 | -3.6 | -2.4 | 0.042 |

To compare the ability of pitchers from different species to compensate weather-related fluctuations of their fluid level, we tested the interaction rainfall × species in a linear mixed-effects model with pitcher ID as a random effect. We found that rain-induced fluid level increase differed between species (likelihood ratio test of rainfall previous day × daily fluid level change: χ^2^=16.7, p=0.005), with *N. hemsleyana* showing the largest and *N. albomarginata* the smallest increase.

### 3.2 Pitcher fluid properties

Pitcher fluid pH, conductivity, K+ concentration, surface tension and viscoelastic properties (relaxation time) all differed significantly between the six *Nepenthes* species studied (Fig. 2, Tables S1, S2; all ANOVAs & Welch-ANOVAs, p<0.001).

The fluids’ conductivity, acidity and K+ concentrations appeared to be correlated. The highest K+ concentrations were found in the species with the most acidic pitcher fluid, *N. rafflesiana* and *N. gracilis*, while the lowest were found in the species with the least acidic fluids (*N. ampullaria* and *N. bicalcarata*) (Fig. 2, Table S1).

The fluids of *N. hemsleyana*, *N. rafflesiana* and *N. gracilis* were the only ones that showed at least briefly (>2 frames at 300 frames per second) a fluid filament when stretched in the extensional rheometer. The fluids of these three species also had a slightly reduced surface tension (Fig.2, Table S1). All species except *N. ampullaria* and *N. bicalcarata* had a significantly lower surface tension than water (72.1 ± 1.3 mN/m, measured *in situ* using our experimental apparatus) (one-sample t-tests, p>0.05).

### 3.3 Effect of pitcher fluid level on prey capture efficiency

We found different effects of pitcher fluid level on insect prey capture efficiency in the six *Nepenthes* species studied (Fig. 3). *N. ampullaria* and *N. bicalcarata* proved to be consistently efficient traps and there was no significant difference in the number of ants captured at low, intermediate, high and full pitcher fluid levels. In *N. albomarginata* and *N. hemsleyana* pitchers, prey capture efficiency was significantly lower when they were completely filled with fluid, as the ants were able to escape more easily across the peristome. Similarly, *N. gracilis* showed a gradual decrease in prey capture efficiency with increasing fluid level. *N. rafflesiana* had its highest prey capture efficiency at intermediate and high pitcher fluid levels, in agreement with Andrew *et al*. (2026).

**Figure 3:**
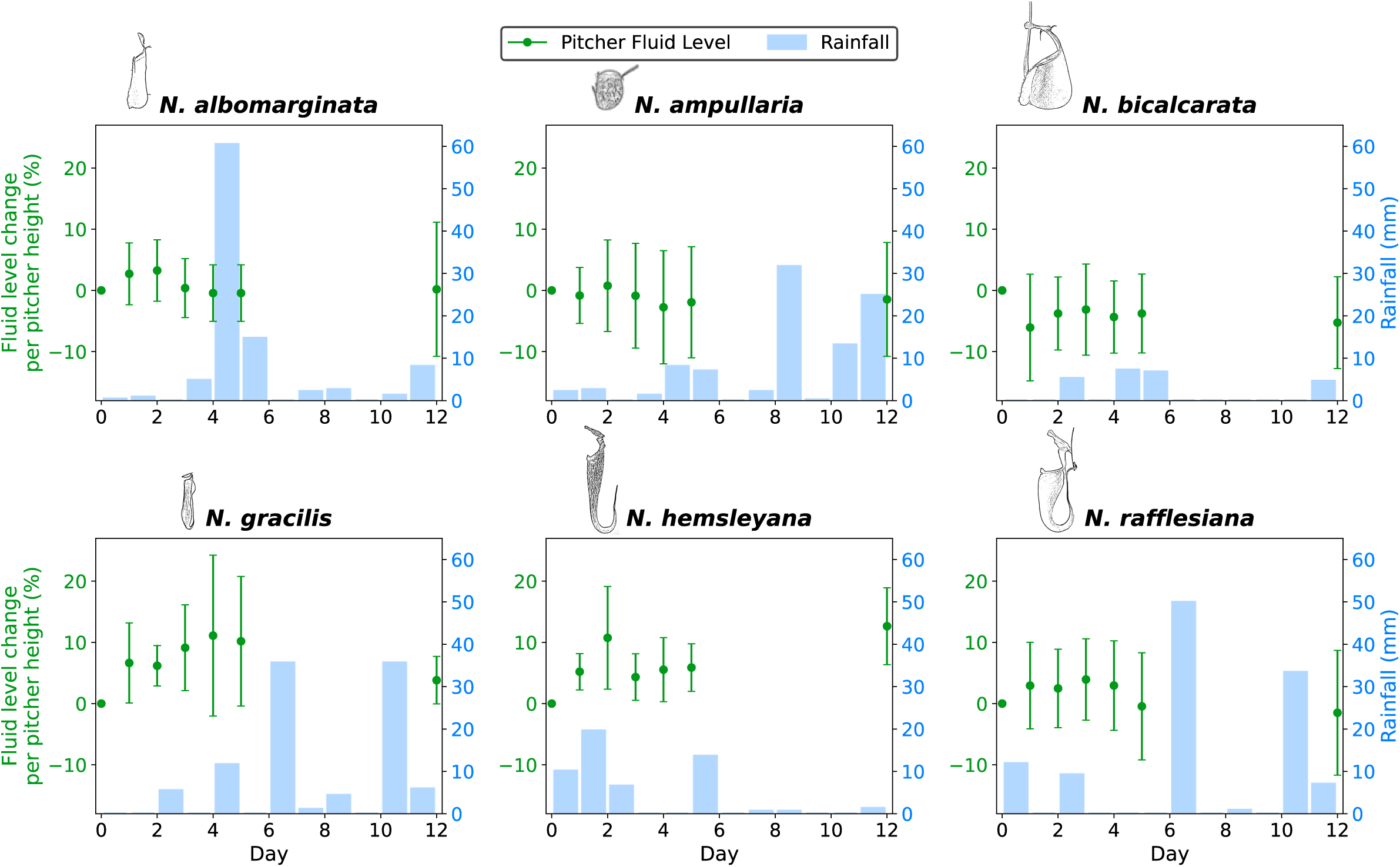
Field measurements of the fluid level change relative to total pitcher height (in %) over a 12-day period for six *Nepenthes* species (green; mean ± standard error) and the rainfall measured at each of the field sites during the experimental period (blue bars). **Alt text:** Graphs showing the daily change in pitcher fluid level compared to the amount of rainfall for each species.

**Figure 4:**
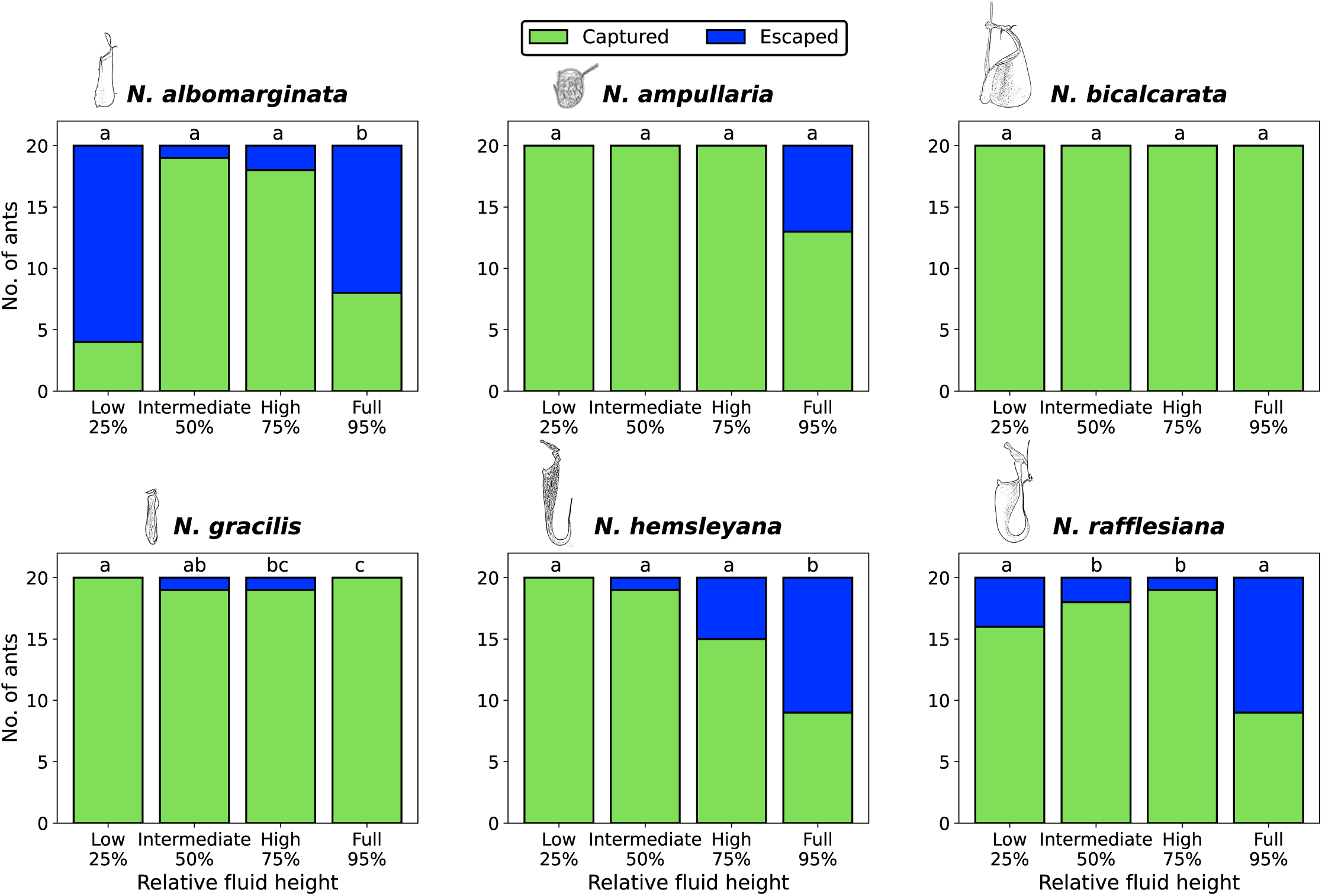
Prey capture efficiency of the pitchers of six *Nepenthes* species at different pitcher fluid levels. One pitcher per species was used; its fluid level was set to one of four levels, if necessary by adding fresh fluid from other pitchers of the same species. The fate of the first 20 *Dolichoderus cuspidatus* ants that slipped and fell into the pitcher fluid was recorded as ‘captured’ by the pitcher (green) or ‘escaped’ (within 5 min, blue). Different letters indicate statistically significant differences between fluid levels according to pairwise Fisher exact tests (p < 0.05), while groups sharing the same letter are not significantly different. **Alt text:** Bar charts showing the number of ants captured and escaped at 4 different pitcher fluid heights in each species.

### 3.4 Response to manipulations of the pitcher fluid volume

When we added pitcher fluid to pitchers so that they were filled to their peristome, all six species showed a significant decrease in fluid level over the 12-day period (Fig. 5; paired t-tests, fluid level on day 0 versus day 12: all species: p<0.01), but they differed significantly in their rate of fluid level decrease (Fig. 9a, ANOVA: F_5,54_=5.7, p=0.001. *N. ampullaria* showed the highest initial rate (relative fluid level change over the first 24 hours of the experiment), while all other species did not differ significantly (Fig. 9a). Only in *N. albomarginata* and *N. ampullaria* did the fluid levels return to near their initial value (paired t-tests, original versus final fluid level: both p > 0.1); fluid levels remained significantly higher in the other species (paired t-tests: all p < 0.01).

**Figure 5:**
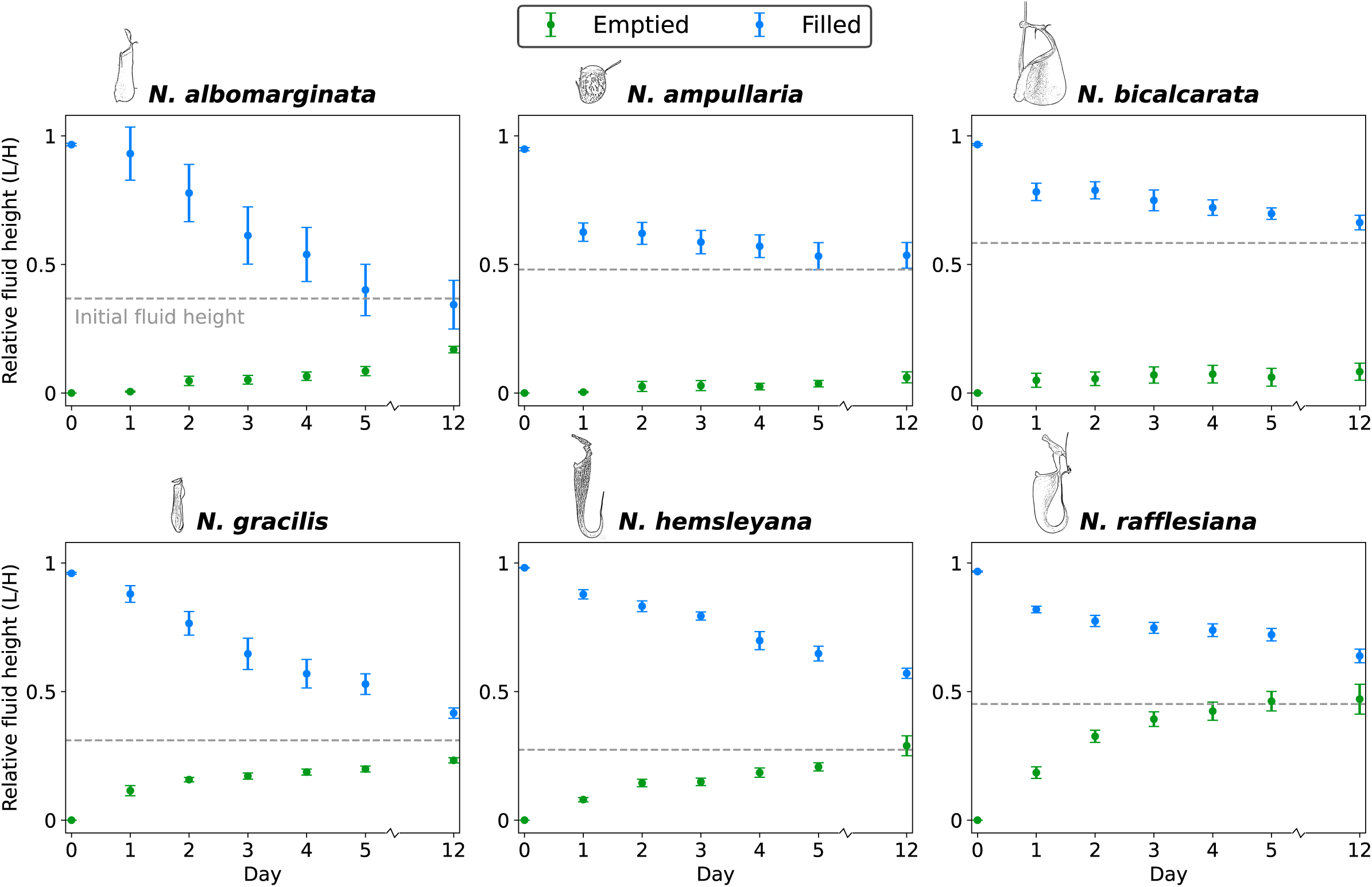
Relative fluid level (fluid level L/ pitcher height H) following volume manipulation (emptying or filling) in six *Nepenthes* species. In ten pitchers of each species, the fluid was removed (green) and ten pitchers were filled to their peristome with fluid from other pitchers of the same species (blue). All values are plotted as mean ± standard error. The grey dashed line marks the average initial relative fluid level for each species. **Alt text:** Graphs showing the daily relative fluid height in pitchers of each species following the pitchers being either emptied of all fluid or filled to the peristome with fluid.

When pitchers were completely emptied of their fluid, all six species showed a significant increase in fluid level over 12 days (Fig. 5; paired t-tests, fluid level on day 0 versus day 12: all p < 0.05), but they differed significantly in their initial rate of fluid replenishment (Welch-ANOVA: F_5,54_=30.3, p<0.001). The highest initial rate of fluid level increase was recorded in *N. rafflesiana* and the lowest in *N. albomarginata* and *N. ampullaria* (Fig. 9b). *N. hemsleyana* and *N. rafflesiana* replenished pitcher fluid so efficiently that within 12 days, fluid levels were not significantly different from the original values (paired t-tests, original versus final fluid level: all p > 0.05).

### 3.5 Response to changes in ionic concentration

When pitcher fluid was replaced with 100 mM KCl up to the original fluid level, pitchers of all species showed a significant fluid level increase within one day (Fig. 6, Table S2). The rate of fluid level changevaried significantly across species (ANOVA: F_5,54_=29.0, p<0.001); it was fastest in *N. rafflesiana* and slowest in *N. albomarginata* (Fig. 9d). Following this increase, fluid levels remained higher than the original fluid level (Table S2). In all species, conductivity decreased significantly over the course of the experiment (Fig. 7, Table S2), and all species but *N. ampullaria* showed a decrease in K^+^ concentration (Fig. S1). These effects indicate dilution of the fluid through water secretion and/or ion absorption. The product of conductivity and volume, a proxy for the total ionic content in the pitcher fluid, decreased significantly in all species, providing strong evidence of ion absorption (Fig. 7, Table S2). The rate of ion absorption did not differ across the species (Fig. 9h).

**Figure 6:**
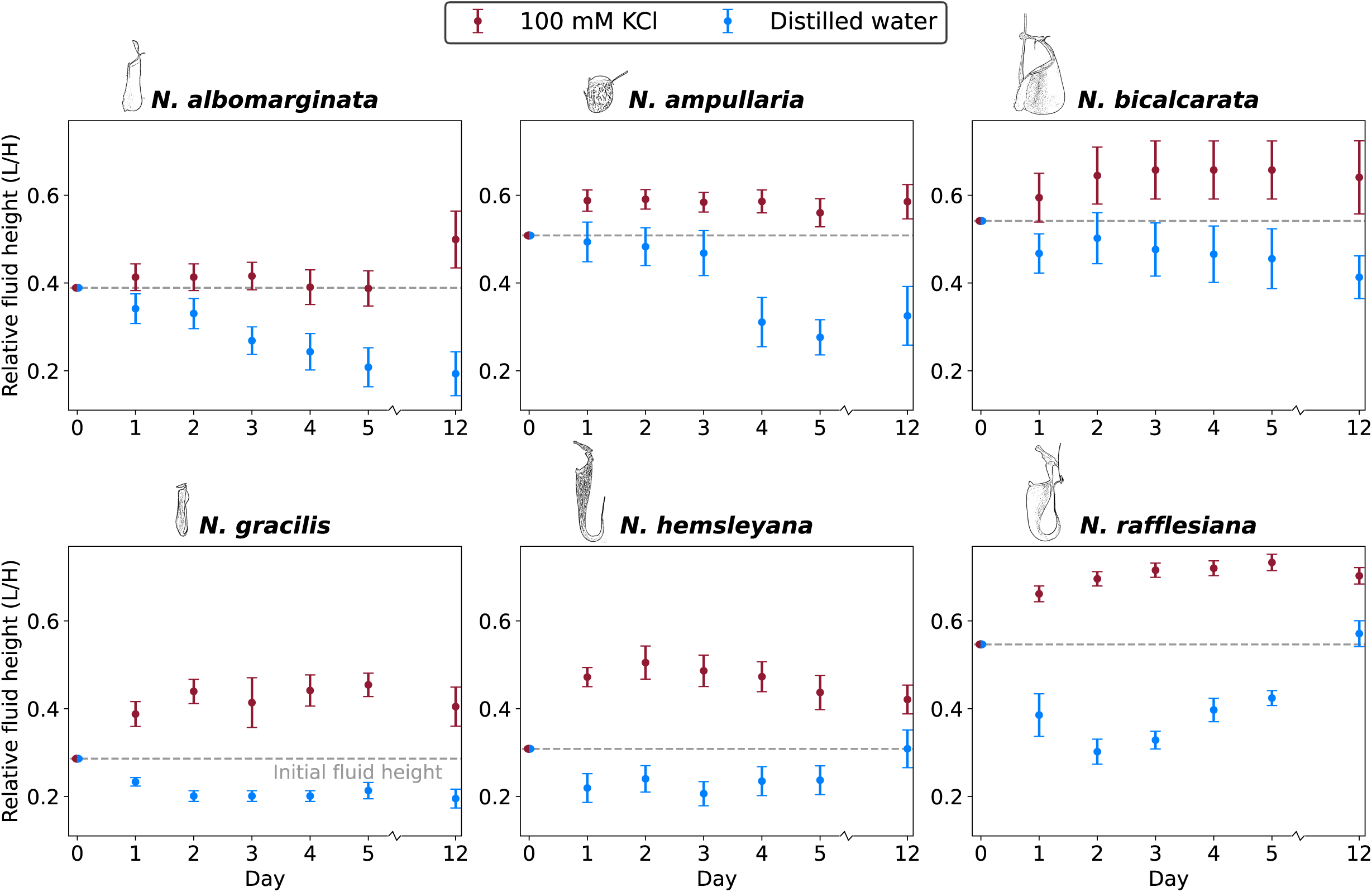
Relative pitcher fluid level (fluid height L/pitcher height H) following fluid replacement with 100 mM KCl or distilled water in six *Nepenthes* species. Values are plotted as mean ± standard error (n=10 pitchers per treatment per species). The grey dashed line marks the average initial relative fluid level for each species. **Alt text:** Graphs showing the daily relative fluid height in pitchers of each species following the pitcher fluid being replaced with either distilled water or 100mM KCl.

**Figure 7:**
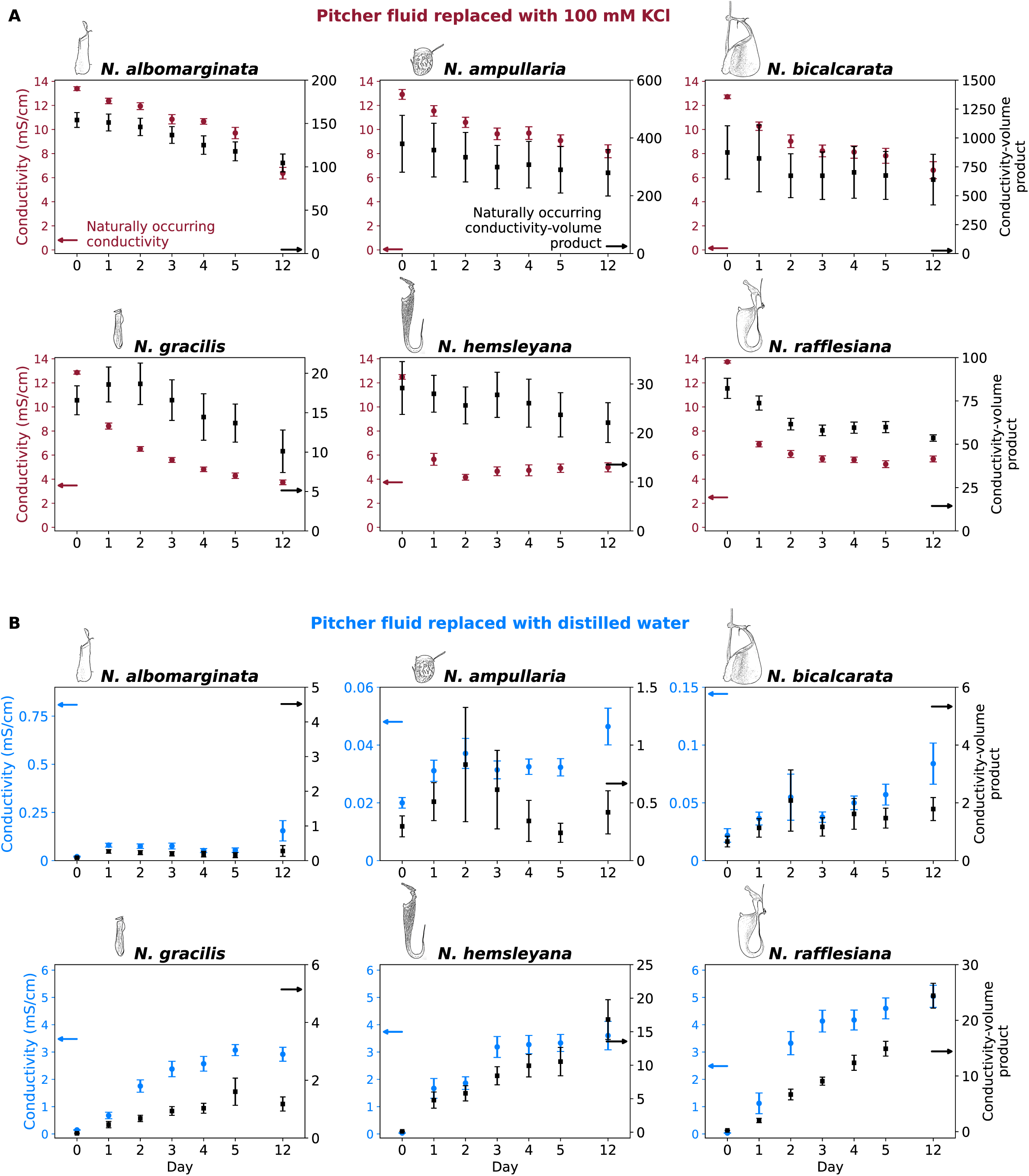
Conductivity (closed circles) and conductivity-volume product normalised by the value measured for each pitcher before the experiment (closed squares) in pitcher fluids from six *Nepenthes* species following replacement with either (A) 100 mM KCl or (B) distilled water. Conductivity-volume product is calculated from the pitcher fluid volume (fig. 5) and serves as a proxy for the total ionic content in the fluid (see also the K+ measurements in Fig. S1). Data are plotted as mean ± standard error (n=10 pitchers per species). Arrows indicate the mean naturally occurring conductivity for the particular species. **Alt text:** Graphs showing the daily conductivity of the pitcher fluid and the product of conductivity and volume measurements of each species following the pitcher fluid being replaced with either distilled water or 100mM KCl. Arrows depict the naturally occurring value of conductivity and conductivity-volume product on the y-axes.

In contrast, when the pitcher fluids of all six species were replaced with distilled water, all species showed a significant increase in conductivity over the course of the experiment, indicating a higher electrolyte concentration due to ion secretion and evaporation/ absorption of fluid (Fig. 7, Table S2). All species with the exception of *N. ampullaria* and *N. bicalcarata* showed evidence of K^+^ secretion, and *N. hemsleyana* and *N. rafflesiana* returned to near their original K^+^ concentration (Table S2). Ion secretion was also evident from the conductivity-volume product, which increased significantly in all species, and with the highest rate in *N. hemsleyana* (Fig. 7, Fig. 9g, Table S2). By secreting ions, pitchers of *N. albomarginata, N. gracilis* and *N. hemsleyana* returned to near their original conductivity, and *N. rafflesiana* even exceeded it (Table S2). Fluid levels in all species decreased, but at significantly different initial rates (ANOVA: F_5,54_=14.3, p<0.001). Fluid levels decreased fastest in *N. rafflesiana* (Fig. 9c). *N. gracilis*, *N. hemsleyana* and *N. rafflesiana* also decreased their fluid levels significantly within one day, suggesting fluid absorption (Fig. 6, Table S2). Fluid levels in *N. albomarginata* and *N. ampullaria* decreased more slowly (Fig. 9c). Following the initial decrease, the pitchers of *N. hemsleyana* and *N. rafflesiana* returned to near their original fluid levels by secreting fluid (Fig. 6).

### 3.6 Response to changes in pitcher fluid pH

Following the addition of 0.1M NaOH to increase the pH of pitcher fluid to pH 11, all species rapidly re-acidified their fluid so that it returned to or even fell below its original pH but the response rate varied across species (Welch-ANOVA: F_5.54_=59.3, p<0.001) (Fig. 8, Fig. 9i, Table S3). *N. gracilis*, *N. hemsleyana* and *N. rafflesiana* showed the fastest rate of re-acidification (in *N. hemsleyana*, the fluid pH dropped from 11.3 to 3.2 in one day), while re-acidification was slowest and took more than 3 days in *N. bicalcarata*.

**Figure 8:**
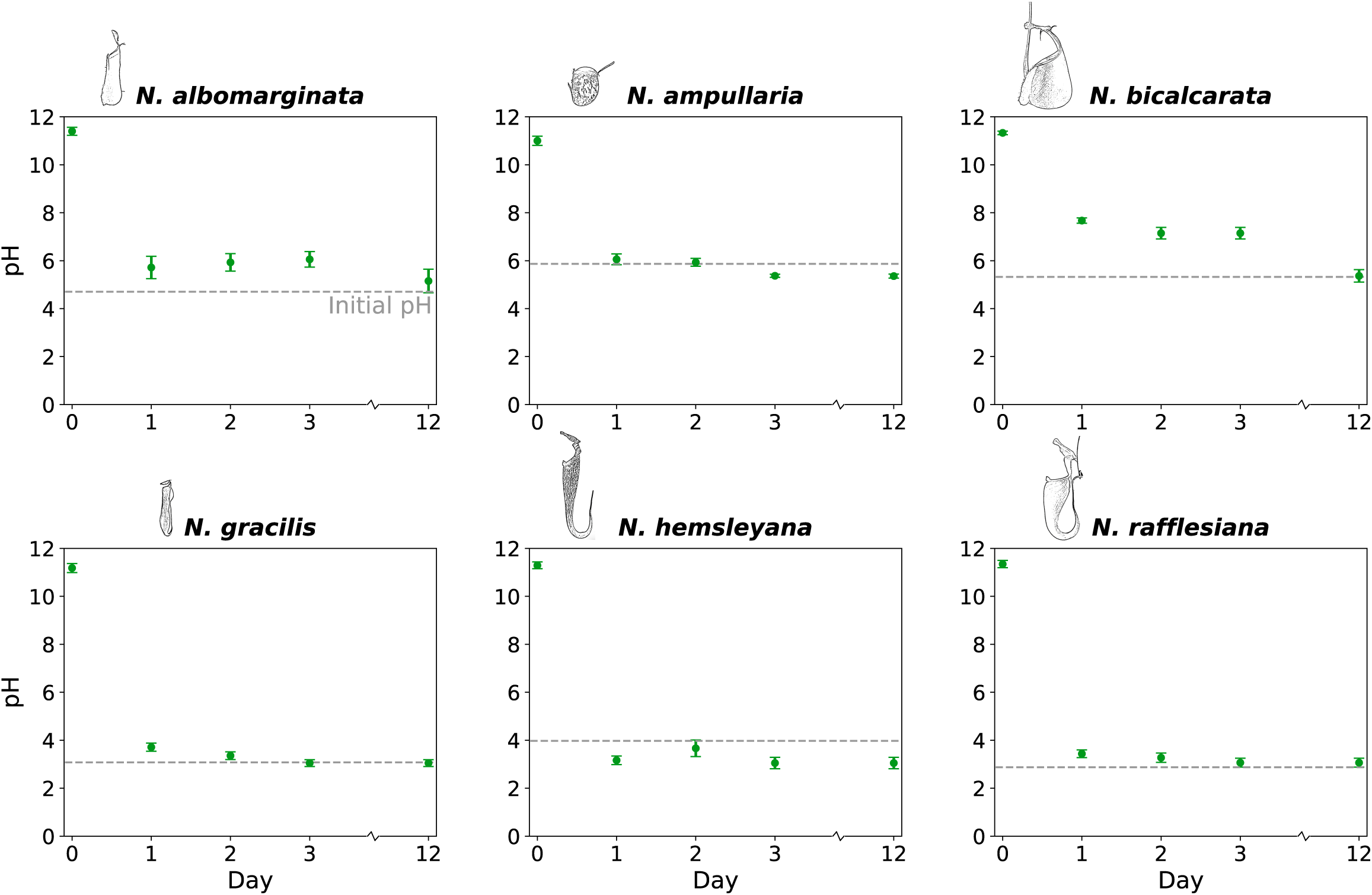
Pitcher fluid pH following the addition of NaOH, raising the pH to 11 in six *Nepenthes* species. pH values are plotted as mean ± standard error (n=10 pitchers per species). The blue dashed line marks the naturally occurring pH for each species. **Alt text:** Graphs showing the daily pH of the fluid in pitchers of each species following the pitcher fluid being doped with alkaline NaOH.

### 3.7 Correlation between physiological rates in the different Nepenthes species

We measured nine different physiological responses of *Nepenthes* pitchers to experimental manipulations:

1. Fluid volume absorption when filled with pitcher fluid
2. Fluid volume secretion when emptied
3. Fluid volume absorption when replaced with distilled water
4. Fluid volume secretion when replaced with high ion concentration
5. Ion absorption (conductivity-volume product decrease) when filled with pitcher fluid
6. Ion secretion (conductivity-volume product increase) when emptied
7. Ion secretion (conductivity-volume product increase) when replaced with distilled water
8. Ion absorption (conductivity-volume product decrease) when replaced with high ion concentration
9. pH drop after alkalisation

To determine the extent to which the average rates of these nine physiological functions measured for each *Nepenthes* species are independent of one another or correlated, we calculated pairwise correlations (Table S4). The correlation matrix shows that all 36 pairwise correlations were positive, and 8 of these correlations were significant (uncorrected p-values <0.05; Table S4).

These positive correlations suggest that all the measured rates, including fluid/electrolyte secretion and absorption, as well as concentration- and volume-dependent fluid regulation, are based on the same or interrelated physiological structures and processes, most likely the activity of the glands on the inner pitcher wall. This activity was highest in *N. rafflesiana* and *N. hemsleyana,* intermediate in *N. gracilis*, and low in *N. ampullaria*, *N. albomarginata* and *N. bicalcarata*.

## Discussion

All *Nepenthes* species investigated in this study showed the ability to regulate their pitcher fluid volume and composition. Pitchers generally remained approximately half-filled with fluid despite environmental fluctuations, in line with previous observations (Hooker, 1874; Clarke, 1997; Andrew *et al*., 2026). This regulation of pitcher fluid volume is likely to increase insect capture rate as intermediate fluid levels maximised prey capture efficiency in most species. Prey capture was significantly reduced when there was either insufficient fluid to drown the insect, or when prey could escape over the peristome. While prey capture in other species was not as strongly dependent on an intermediate fluid level as in *N. rafflesiana* (Andrew *et al*., 2026), the fluid level responses we measured suggested that all *Nepenthes* species prevent their pitchers from becoming empty or flooded. to maintain sufficient prey capture. Apart from the benefit it brings to prey capture, ensuring a sufficient volume of fluid in the pitcher is also crucial for the enzymatic digestion of prey and for the survival of the pitcher infauna. This community of dipteran larvae, mites and bacteria helps with the breakdown of prey and makes nutrients available to the plant (Lam *et al*., 2017; Leong *et al*., 2018; Lam *et al*., 2019a).

All the *Nepenthes* species were able to absorb and secrete fluid. Variation of the conductivity-volume product and direct measurements of K+ concentrations demonstrate that all the species are able to absorb ions, and all species except *N. ampullaria* and *N. bicalcarata* showed evidence for ion secretion. All species demonstrated the ability to rapidly secrete protons to maintain the acidity of the pitcher fluid. Our findings indicate that the control of pitcher fluid level can be triggered by both changes in the volume and concentration of the pitcher fluid, as suggested for *N. rafflesiana* (Andrew *et al*., 2026). All species secreted fluid when pitchers were emptied and all species except *N. bicalcarata* absorbed fluid when pitcher fluid volume was increased. When pitcher fluids were experimentally concentrated, all species responded by secreting fluid and absorbing ions, and all pitchers responded by absorbing fluid and secreting ions when the concentration was experimentally reduced.

Despite the presence of fluid level regulation mechanisms across all the species investigated, the intensity of this control varied significantly. The regulation of pitcher fluid we observed consists of several potentially distinct processes, including concentration and volume-dependent fluid secretion/absorption and ion secretion/absorption, as well as proton secretion. We found that the rates of these processes were positively correlated across the different species (Table S4), indicating that these pitcher fluid regulation mechanisms are linked. A likely explanation is that all the different pitcher fluid regulation processes are driven by the activity of the glandular epithelium lining the inner pitcher wall. Pitchers use these glands to secrete and absorb fluid and ions (Morrissey, 1955; Owen and Lennon, 1999; Moran *et al*., 2010; Adlassnig *et al*., 2011) and our findings indicate that they employ them to regulate their fluid. The differences across species in the intensity of fluid regulation may be explained by the size, function and activity of these glands (Owen *et al*., 2014), and the location of these glandular zones within the pitcher. Species with weaker regulatory capacity such as *N. albomarginata, N. ampullaria* and *N. bicalcarata* may mainly rely on passive fluid level control via evaporation and osmotic water transport.

The regulation of fluid volume in *Nepenthes* pitchers appears to be faster and more active than the fluid regulation in other, convergently evolved pitcher plant genera. In *Sarracenia* pitchers, fluid absorption and secretion by the pitcher was found to be negligible and volumes passively fluctuated along with the rainfall (Kingsolver, 1979; Kingsolver, 1981; Beaver, 1983); in *Cephalotus* pitcher plants seasonally changing fluid volumes have been reported (Clarke, 1988; Adlassnig *et al*., 2011), but the details of its fluid control are still unclear. Fluctuations of open fluid volumes are not limited to carnivorous plants. Fluid level fluctuations due to rainfall can have negative effects on inquiline communities in *Bromeliad* water tanks (Jabiol *et al*., 2009; Romero *et al*., 2020) and the fluid held in nectaries is also influenced by weather conditions and appears to be dynamically regulated (Nepi and Stpiczynska, 2008; Heil, 2011).

The substantial variation between *Nepenthes* species in the intensity of their regulation of pitcher fluid volume and concentration may be explained by several factors, including phylogeny, morphological traits, ecology and life history:

### Phylogenetic relatedness of study species

*Nepenthes* pitcher plants have undergone multiple adaptive radiations, resulting in a species-rich and highly diverse group of carnivorous plants (Meimberg *et al*., 2006; Murphy *et al*., 2020; Scharmann *et al*., 2021). According to the latest available *Nepenthes* phylogeny, all the species used in this study except *N. albomarginata* belong to the same clade, which comprises many widespread, lowland taxa (Murphy *et al*., 2020). Within this clade, *N. rafflesiana* and *N. hemsleyana* are closely related sister species, and both exhibit strong fluid regulation (Fig. 1, Fig. 9). Similarly, *N. bicalcarata* and *N. ampullaria* are closely related, and performed only weak fluid regulation (Fig. 1, Fig. 9). The similar levels of fluid regulation in these species pairs could also have evolved independently, and the number of species investigated here prevents more robust conclusions on the evolutionary history of these traits.

**Figure 9:**
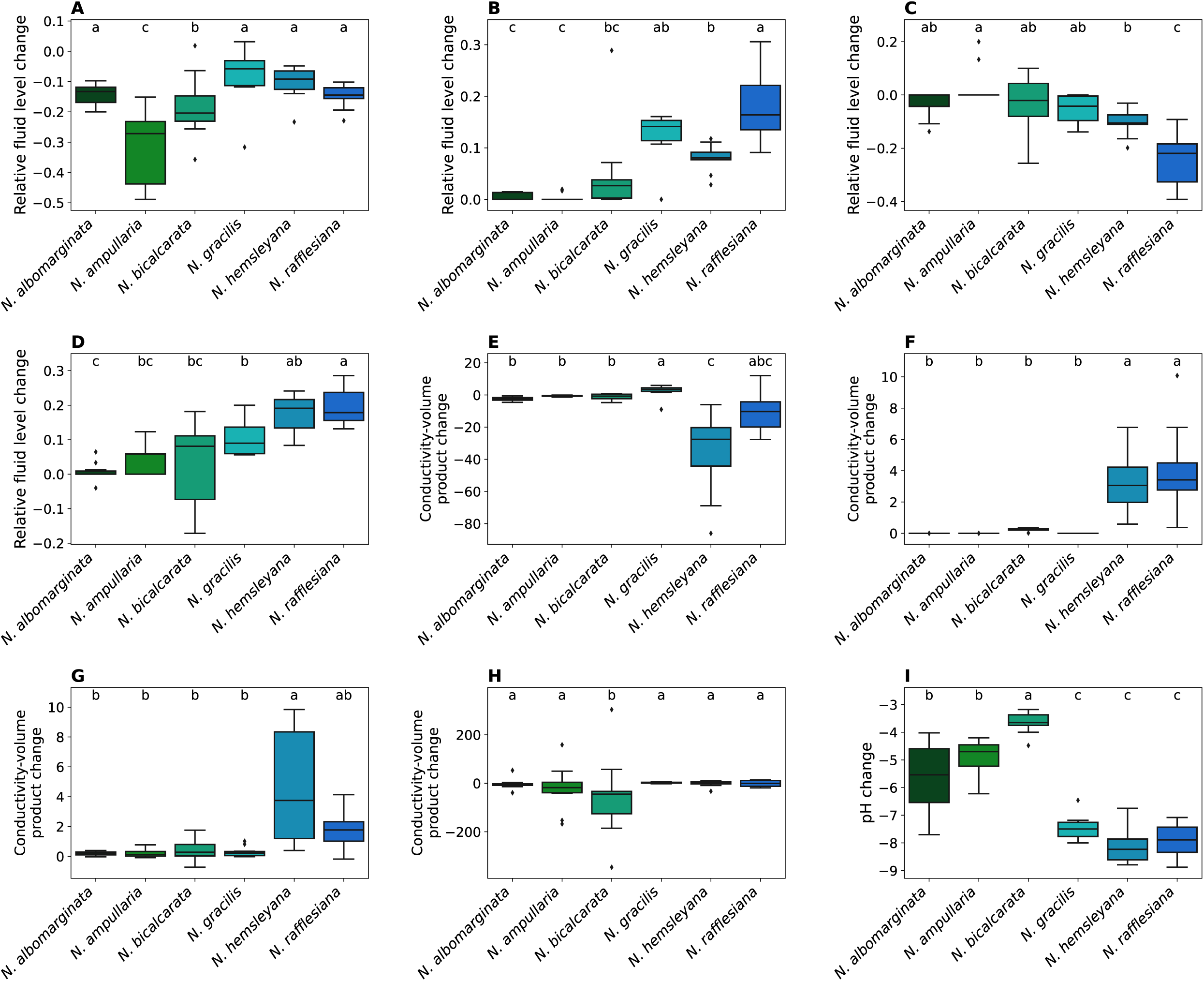
The rate of regulation of pitcher fluid parameters, measured as the change over the first 24 hours following experimental manipulation for all species and all experimental conditions: (a) fluid level change after filling with pitcher fluid, (b) fluid level change after emptying pitcher, (c) fluid level change after pitcher fluid replaced with distilled water, (d) fluid level change after pitcher fluid replaced with 100mM KCl, (e) change in ion content after filling with pitcher fluid, (f) change in ion content after emptying pitcher fluid, (g) change in ion content after pitcher fluid replaced with distilled water, (h) change in ion content after pitcher fluid replaced with 100mM KCl, (i) pH change after NaOH added to pitcher fluid to bring pH to 11. Letters a-c represent the statistical significance of differences between species, based on ANOVA (A, C & D) or Welch-ANOVA (B & E-I) followed by Tukey’s HSD (A, C & D) or Games-Howell (B & E-I) post-hoc tests. **Alt text:** Boxplots comparing the rates of change of the fluid volume and conductivity-volume product in our experiments for each of the six *Nepenthes* species with letters denoting the result of pairwise comparisons.

### Pitcher morphology and trapping mechanism

Differences in the pitchers’ morphology and trapping mechanisms may explain why fluid regulation is more pronounced in some species than in others. Our study species can be broadly classified into peristome-, wax- or fluid-based trapping according to the capture strategies proposed by Moran *et al*. (2013). *N. ampullaria* and *N. bicalcarata* have wide and steep peristomes (Bauer *et al*., 2012a; Moulton *et al*., 2023) and may therefore be considered peristome-trapping species. These species retained prey efficiently regardless of fluid level. It is therefore likely that regulating the fluid level is less important for these species, consistent with the observed lower intensity of their fluid regulation. *N. albomarginata*, *N. gracilis* and *N. hemsleyana* pitchers have slippery wax crystals on their inner wall and may be considered wax-trapping species (Moran *et al*., 2013). In these species, insect capture efficiency strongly declined at high fluid levels (but less so at low fluid levels), probably because the submerged waxy inner surface is ineffective. Lastly, *N. rafflesiana* and *N. hemsleyana* have a sticky pitcher fluid that mediates prey capture (Gaume *et al*., 2007; Bauer *et al*., 2011) and their trapping mechanism may therefore be considered fluid-based (Moran *et al*., 2013). For both species, we observed a decline in capture efficiency for both low and very high fluid levels. Low fluid volumes can have negative effects on prey capture, as many ants that do not land in the pitcher fluid can climb the pitcher walls and escape (Andrew *et al*., 2026).

### Ecology and Life History

#### Habitat

The six *Nepenthes* species used in this study grow in close proximity in Brunei, Borneo, and thus experience similar weather conditions. Nevertheless, species exhibited varying intensities of fluid regulation. Fluid levels in *N. rafflesiana* and *N. hemsleyana* strongly increased with rainfall. Experiments on these species occurred at the same, exposed, field site. *N. gracilis*, also found at this site, did not experience the same degree of fluid level fluctuation, possibly due to its more horizontal lid, which shelters the pitcher more (Chomicki *et al*., 2024). The remaining three species occurred in more sheltered understorey habitats, where we recorded no significant rain-induced fluctuations of fluid level. Reduced fluid volume fluctuations in more sheltered locations have also been recorded in *Sarracenia* and *Cephalotus* pitcher plants (Kingsolver, 1981; Clarke, 1988).

#### Pitcher Lifespan and Construction Costs

A further factor that could explain the varying intensities of fluid regulation is the lifespan of the pitchers, which ranges from a few weeks to 2 ½ years in our study species (Osunkoya *et al*., 2007; Osunkoya *et al*., 2008; Moran and Clarke, 2010; Thornham *et al*., 2012). It has been suggested that a shorter pitcher lifespan is correlated with a higher physiological activity (Osunkoya *et al*., 2008) and potentially a higher investment in efficient prey capture. This is consistent with the fast regulatory responses we observed in the species with the shortest-lived pitchers (*N. rafflesiana* and *N. hemsleyana*) and the slow regulatory responses in *N. bicalcarata*, which has the longest-lived pitchers. However, longer pitcher lifespans are associated with higher construction costs (Osunkoya et al., 2008), which may demand a higher nutrient return from each pitcher. Our findings suggest that pitchers with a longer lifespan, despite their higher construction costs, allow pitcher plants to reduce their investment in fluid regulation.

#### Nutrient acquisition strategies

*Nepenthes* species differ markedly in their nutrient acquisition strategies, and this variation may influence their reliance on pitcher fluid regulation. While many taxa primarily derive nutrients from insect capture, some exhibit unique feeding strategies or engage in symbioses that boost nutrient acquisition in alternative ways. For example, *N. ampullaria* captures falling leaf litter and detritus and has very little reliance on live insect capture, consistent with the weak fluid regulatory responses we observed (Cresswell, 1998; Moran *et al*., 2003; Pavlovic *et al*., 2011; Gilbert *et al*., 2022). In *N. hemsleyana*, Hardwicke’s Woolly bats roost in the pitchers and their excrement contributes an estimated 33.8 % of the pitcher’s total nitrogen (Grafe *et al*., 2011). Apart from the slender morphology of its pitchers with the sudden taper, the low fluid levels may facilitate bat roosting (Schöner *et al*., 2013; Lim *et al*., 2014; Schöner *et al*., 2015a) by providing a safe dry zone above the fluid. To ensure good conditions for the bats, it is important for *N. hemsleyana* to efficiently maintain its fluid at a low level, consistent with its rapid response to fluid volume and composition changes. The pitchers of *N. gracilis* and *N. rafflesiana* are regularly inhabited by crab spiders (*Misumenops nepenthicola* and *Thomisus nepenthiphilus*). These spiders ambush struggling prey, preventing it from escaping and depositing waste products that benefit overall nutrient intake (Lam *et al*., 2019b; Karl and Bauer, 2020). Karl and Bauer (2020) observed spiders promptly abandoning pitchers when the fluid level became too high or too low. In order to facilitate this mutualistic interaction, the two *Nepenthes* species need to maintain an intermediate fluid level, consistent with the efficient pitcher fluid regulation we observed.

## Conclusion

We demonstrate that many *Nepenthes* species have the ability to regulate their pitcher fluid, and that the intensity of this regulation varies markedly, consistent with each species’ pitcher morphology, nutrient acquisition strategy and other ecological traits. Our findings show how pitcher plants with diverse ecological strategies are affected by fluctuating weather conditions and how they can compensate for these effects through regulation of their pitcher fluid.

## Acknowledgements

The authors would like to thank Universiti Brunei Darussalam for their logistical support during our 2024 field season and the Brunei Forestry Department for issuing forest entry permits. Our extended thanks go to the Tinggal, Rahaman and Engin families for their warm reception and assistance during our stay in Brunei and to Ritabrata Chowdhury for his support during field data collection.

## Author Contributions

C.N.S.A: conceptualisation, methodology, data collection, data analysis, writing-initial draft, reviewing and editing. J.Y.B and I.J.H: data collection, writing-reviewing and editing. F.M and T.U.G: reviewing and editing. W.F: supervision, conceptualisation, methodology, writing-initial draft, reviewing and editing.

## References

Adlassnig W, Peroutka M, Lendl T. 2011. Traps of carnivorous pitcher plants as a habitat: composition of the fluid, biodiversity and mutualistic activities. Annals of Botany 107: 181–194. DOI: 10.1093/aob/mcq238

Andrew C, Bu J, Kelly N, et al. Federle W. 2026. An insect trap adjusting to weather conditions: Nepenthes rafflesiana plants control the fluid level in their pitchers to maximise prey capture. Annals of Botany 137: 833–846. DOI: 10.1093/aob/mcaf294

Bartholomeus RP, Witte J-PM, Bodegom PMv, Dam JCv, Aerts R. 2011. Climate change threatens endangered plant species by stronger and interacting water-related stress. Journal of Geophysical Research 116: G04023. DOI: 10.1029/2011JG001693

Bauer U, Bohn H, Federle W. 2008. Harmless nectar source or deadly trap: Nepenthes pitchers are activated by rain, condensation and nectar. Proceedings of the Royal Society B 275: 259–265. DOI: 10.1098/rspb.2007.1402

Bauer U, Clemente C, Renner T, Federle W. 2012a. Form follows function: morphological diversification and alternative trapping strategies in carnivorous Nepenthes pitcher plants. Journal of Evolutionary Biology 25: 90–102. DOI: 10.1111/j.1420-9101.2011.02406.x

Bauer U, Federle W, Seidel H, Grafe T, Ioannou C. 2015. How to catch more prey with less effective traps: explaining the evolution of temporarily inactive traps in carnivorous pitcher plants. Proceedings of the Royal Society B 282: 20142675. DOI: 10.1098/rspb.2014.2675

Bauer U, Giusto BD, Skepper J, Grafe TU, Federle W, Ollerton J. 2012b. With a flick of the lid: A novel trapping mechanism in Nepenthes gracilis pitcher plants. PLos One 7: e38951. DOI: 10.1371/journal.pone.0038951

Bauer U, Grafe TU, Federle W. 2011. Evidence for alternative trapping strategies in two forms of the pitcher plant, Nepenthes rafflesiana. Journal of Experimental Botany 62: 3683–3692. DOI: 10.1093/jxb/err082

Bauer U, Willmes C, Federle W. 2009. Effect of pitcher age on trapping efficiency and natural prey capture in carnivorous Nepenthes rafflesiana plants. Annals of Botany 103: 1219–1226. DOI: 10.1093/aob/mcp065

Beaver R. 1983. The communities living in Nepenthes pitcher plants: fauna and food webs. In: Phytotelmata: Terrestrial Plants as Hosts for Aquatic Insect Communities. (ed. Frank, JH, Lounibos, LP). Medford, New Jersey: Plexus Publishing Inc., 129–159.

Bennett K, Ellison A. 2009. Nectar, not colour, may lure insects to their death. Biology Letters 5: 469–472. DOI: 10.1098/rsbl.2009.0161

Bohn H, Federle W. 2004. Insect aquaplaning: Nepenthes pitcher plants capture prey with the peristome, a fully wettable water-lubricated anisotropic surface. PNAS 101: 14138–14143. DOI: 10.1073/pnas.0405885101

Bonhomme V, Pelloux-Prayer H, Jousselin E, Forterre Y, Labat JJ, Gaume L. 2011. Slippery or sticky? Functional diversity in the trapping strategy of Nepenthes carnivorous plants. New Phytol 191: 545–554. DOI: 10.1111/j.1469-8137.2011.03696.x

Chin L, Moran J, Clarke C. 2009. Trap geometry in three giant montane pitcher plant species from Borneo is a function of tree shrew body size. New Phytologist 186: 461–470. DOI: 10.1111/j.1469-8137.2009.03166.x

Chomicki G, Burin G, Busta L, et al. Bauer U. 2024. Convergence in carnivorous pitcher plants reveals a mechanism for composite trait evolution. Science 383: 108–113. DOI: 10.1126/science.ade0529

Clarke C. 1997. Nepenthes of Borneo. Kota Kinabalu: Natural History Publications.

Clarke C, Bauer U, Lee C, Tuen A, Rembold K, Moran J. 2009. Tree shrew lavatories: a novel nitrogen sequestration strategy in a tropical pitcher plant. Biology Letters 5: 632–635. DOI: 10.1098/rsbl.2009.0311

Clarke C, Kitching R. 1993. The metazoan food webs from six Bornean Nepenthes species. Ecological Entomology 18: 7–16. DOI: 10.1111/j.1365-2311.1993.tb01074.x

Clarke S. 1988. Seasonal growth and mortality of the pitchers of the Albany pitcher plant, *Cephalotus follicularis* Labill. Australian Journal of Botany 36: 643–653. DOI: 10.1071/BT9880643

Collett C, Ardron A, Bauer U, et al. Pratt L. 2015. A portable extensional rheometer for measuring the viscoelasticity of pitcher plant and other sticky liquids in the field. Plant Methods 11: 16. DOI: 10.1186/s13007-015-0059-5

Cresswell J. 1998. Morphological correlates of necromass accumulation in the traps of an Eastern tropical pitcher plant, Nepenthes ampullaria Jack, and observations on the pitcher infauna and its reconstitution following experimental removal. Oecologia 113: 383–390. DOI: 10.1007/s004420050390

Di Giusto B, Grosbois V, Fargeas E, Marshall D, Gaume L. 2008. Contribution of pitcher fragrance and fluid viscosity to high prey diversity in a Nepenthes carnivorous plant from Borneo. Journal of Biosciences 33: 121–136. DOI: 10.1007/s12038-008-0028-5

Ellison A, Adamec L. 2018. Introduction: what is a carnivorous plant? In: Carnivorous Plants: Physiology, ecology and evolution (ed. Ellison, A, Adamec, L). Oxford: Oxford University Press, 408–410. DOI: 10.1093/oso/9780198779841.003.0029

Frank J. 1983. Bromeliad phytotelmata and their biota, especially mosquitoes. In: Phytotelmata: Terrestrial plants as hosts for aquatic insect communities (ed. Frank, JH, Lounibos, LP). Medford, New Jersey: Plexus Publishing Inc.

Gaume L, Bazile V, Bousses P, Moguedec GL, Marshall D. 2019. The biotic and abiotic drivers of ‘living’ diversity in the deadly traps of Nepenthes pitcher plants. Biodiversity and Conservation 28: 345–362. DOI: 10.1007/s10531-018-1658-z

Gaume L, Forterre Y, Lynn D. 2007. A viscoelastic deadly fluid in carnivorous pitcher plants. PLos One 2: e1185. DOI: 10.1371/journal.pone.0001185

Gaume L, Perret P, Gorb E, Gorb S, Labat J, Rowe N. 2004. How do plant waxes cause flies to slide? Experimental tests of wax-based trapping mechanisms in three pitfall carnivorous plants. Arthropod Structure and Development 33: 103–111. DOI: 10.1016/j.asd.2003.11.005

Gerlach G. 2011. The genus Coryanthes: A paradigm in ecology. Lankesteriana 11: 253–264. DOI: 10.15517/lank.v11i3.18280

Gilbert K, Goldsborough T, Lam W, Leong F, Pierce N. 2022. A semi-detritivorous pitcher plant, Nepenthes ampullaria diverges in its regulation of pitcher fluid properties. Journal of Plant Interactions 17: 956–966. DOI: 10.1080/17429145.2022.2123567

Gorb E, Haas K, Henrich A, Enders S, Barbakadze N, Gorb S. 2005. Composite structure of the crystalline epicuticular wax layer of the slippery zone in the pitchers of the carnivorous plant Nepenthes alata and its effect on insect attachment. Journal of Evolutionary Biology 208: 4651–4662. DOI: 10.1242/jeb.01939

Grafe T, Kohout R. 2013. A new case of ants nesting in Nepenthes pitcher plants. Ecotropica 19: 77–80.

Grafe T, Schöner C, Kerth G, Junaidi A, Schöner M. 2011. A novel resource-service mutualism between bats and pitcher plants. Biology Letters 7: 436–439. DOI: 10.1098/rsbl.2010.1141

Gray L, Clarke C, Wint G, Moran J. 2017. Potential effects of climate change on members of the Palaeotropical pitcher plant family Nepenthaceae. PLos One 12: e0183132. DOI: 10.1371/journal.pone.0183132

Heil M. 2011. Nectar: generation, regulation and ecological functions. Trends in Plant Science 16: 191–200. DOI: 10.1016/j.tplants.2011.01.003

Hooker J. 1874. The carnivorous habits of plants. Nature 10: 366–372. DOI: 10.1038/010366a0

Huang E, Skoufis A, Denning T, et al. Berry J. 2021. OpenDrop: Open-source software for pendant drop tensiometry & contact angle measurements. Journal of Open Source Software 6: 2604. DOI: 10.21105/joss.02604

Jabiol J, Corbara B, Dejean A, Cereghino R. 2009. Structure of aquatic insect communities in tank-bromeliads in a East-Amazonian rainforest in French Guiana. Forest Ecology and Management 257: 351–360. DOI: 10.1016/j.foreco.2008.09.010

Juniper B, Robins R, Joel D. 1989. The Carnivorous Plants. London: Academic Press.

Kang V, Isermann H, Sharma S, Wilson D, Federle W. 2021. How a sticky fluid facilitates prey retention in a carnivorous pitcher plant (Nepenthes rafflesiana). Acta Biomaterialia 128: 357–369. DOI: 10.1016/j.actbio.2021.04.002

Karl I, Bauer U. 2020. Inside the trap: Biology and behavior of the pitcher-dwelling crab spider, Misumenops nepenthicola. *Plants, People*, Planet 2: 290–293. DOI: 10.1002/ppp3.10104

Kingsolver J. 1979. Thermal and hydric aspects of environmental heterogeneity in the pitcher plant mosquito. Ecological Monographs 49: 357–376. DOI: 10.2307/1942468

Kingsolver J. 1981. The effect of environmental uncertainty on morphological design and fluid balance in Sarracenia purpurea. Oecologia 48: 364–370. DOI: 10.1007/BF00346496

Kitching R. 2000. Food Webs and Container Habitats : The Natural History and Ecology of Phytotelmata. Cambridge, UK: Cambridge University Press.

Lam W, Chong K, Anand G, Tan H. 2017. Dipteran larvae and microbes facilitate nutrient sequestration in the Nepenthes gracilis pitcher plant host. Biology Letters 13: 20160928. DOI: 10.1098/rsbl.2016.0928

Lam W, Chou Y, Leong F, Tan H. 2019a. Inquiline predator increases nutrient-cycling efficiency of Nepenthes rafflesiana pitchers. Biology Letters 15: 20190691. DOI: 10.1098/rsbl.2019.0691

Lam W, Ling J, Lum T, Tan H. 2020. Ecology and natural history of swimming pitcher mites (Creutzeria spp., Histiostomatidae) from the traps of Nepenthes pitcher plants. Journal of Zoology 310: 1–9. DOI: 10.1111/jzo.12727

Lam W, Tan H, Dussutour A. 2019b. The crab spider–pitcher plant relationship is a nutritional mutualism that is dependent on prey-resource quality. Journal of Animal Ecology 88: 102–113. DOI: 10.1111/1365-2656.12915

Lenz A, Bauer U. 2022. Pitcher geometry facilitates extrinsically powered ‘springboard trapping’ in carnivorous Nepenthes gracilis pitcher plants. Biology Letters 18: 20220106. DOI: 0.1098/rsbl.2022.0106

Leong F, Lam W, Tan H. 2018. A dipteran larva–pitcher plant digestive mutualism is dependent on prey resource digestibility. Oecologia 188: 813–820. DOI: 10.1007/s00442-018-4258-4

Lim Y, Schöner C, Schöner M, et al. Grafe T. 2014. How a pitcher plant facilitates roosting of mutualistic woolly bats. Evolutionary Ecology Research 16: 581–591.

McPherson S. 2023. Nepenthes-The Tropical Pitcher Plants. Poole, Dorset, England: Redfern Natural History Productions.

Meimberg H, Thalhammer S, Brachmann A, Heubl G. 2006. Comparative analysis of a translocated copy of the trnK intron in carnivorous family Nepenthaceae. Molecular Phylogenetics and Evolution 39: 478–490. DOI: 10.1016/j.ympev.2005.11.023

Merbach M, Merbach D, Maschwitz U, Booth W, Fiala B, Zizka G. 2002. Mass march of termites into the deadly trap. Nature 415: 36–37. DOI: 10.1038/415036a

Moran J. 1996. Pitcher dimorphism, prey composition and the mechanisms of prey attraction in the pitcher plant Nepenthes rafflesiana in Borneo. Journal of Ecology 84: 515–525. DOI: 10.2307/2261474

Moran J, Clarke C. 2010. The carnivorous syndrome in Nepenthes pitcher plants: current state of knowledge and potential future directions. Plant Signaling & Behaviour 5: 644–648. DOI: 10.4161/psb.5.6.11238

Moran J, Clarke C, Hawkins B. 2003. From carnivore to detritivore? Isotopic evidence for leaf litter utilization by the tropical pitcher plant Nepenthes ampullaria. . International Journal of Plant Sciences 164: 635–639. DOI: 10.1086/375422

Moran J, Gray L, Clarke C, Chin L. 2013. Capture mechanism in Palaeotropical pitcher plants (Nepenthaceae) is constrained by climate. Annals of Botany 112: 1279–1291. DOI: 10.1093/aob/mct195

Moran J, Hawkins B, Gowen B, Robbins S. 2010. Ion fluxes across the pitcher walls of three Bornean Nepenthes pitcher plant species: flux rates and gland distribution patterns reflect nitrogen sequestration strategies. Journal of Experimental Botany 61: 1365–1374. DOI: 10.1093/jxb/erq004

Moran J, Merbach M, Livingston N, Clarke C, Booth W. 2001. Termite prey specialization in the pitcher plant Nepenthes albomarginata-evidence from stable isotope analysis. Annals of Botany 88: 307–311. DOI: 10.1006/anbo.2001.1460

Morrissey S. 1955. Chloride ions in the secretion of the pitcher plant. Nature 176: 1220–1221.

Moulton D, Oliveri H, Goriely A, Thorogood C. 2023. Mechanics reveals the role of peristome geometry in prey capture in carnivorous pitcher plants (Nepenthes). PNAS 120: e2306268120. DOI: 10.1073/pnas.2306268120

Murphy B, Forest F, Barraclough T, et al. Cheek M. 2020. A phylogenomic analysis of Nepenthes (Nepenthaceae). Molecular Phylogenetics and Evolution 144: 106668. DOI: 10.1016/j.ympev.2019.106668

Nepi M, Stpiczynska M. 2008. Do plants dynamically regulate nectar features through sugar sensing? Plant Signaling & Behaviour 10: 874–876. DOI: 10.4161/psb.3.10.6228

Osunkoya O, Daud S, Di Giusto B, Wimmer F, Holige T. 2007. Construction costs and physico-chemical properties of the assimilatory organs of Nepenthes species in Northern Borneo. Annals of Botany 99: 895–906. DOI: 10.1093/aob/mcm023

Osunkoya O, Daud S, Wimmer F. 2008. Longevity, lignin content and construction cost of the assimilatory organs of Nepenthes species. Annals of Botany 102: 845–853. DOI: 10.1093/aob/mcn162

Owen T, Carini A, Sutherland L, Hass C, Gabow K. 2014. Structure and development of the attractive and digestive glands in the carnivorous pitcher plant Nepenthes alata. Microscopy and Microanalyses 20. DOI: 10.1017/S1431927614008186

Owen T, Lennon K. 1999. Structure and Development of the Pitchers from the Carnivorous Plant Nepenthes Alata (Nepenthaceae). American Journal of Botany 86: 1382–1390. DOI: 10.2307/2656921

Pavlovic A, Slovakova L, Santrucek J. 2011. Nutritional benefit from leaf litter utilization in the pitcher plant Nepenthes ampullaria. Plant, Cell and Environment 34: 1865–1873. DOI: 10.1111/j.1365-3040.2011.02382.x

Pickering B, Duff T, Baillie C, Cawson J. 2021. Darker, cooler, wetter: forest understories influence surface fuel moisture. Agricultural and Forest Meteorology 300: 108311. DOI: 10.1016/j.agrformet.2020.108311

Riedel M, Eichner A, Meimberg H, Jetter R. 2007. Chemical composition of epicuticular wax crystals on the slippery zone of pitchers of five Nepenthes species and hybrids. Planta 225: 1517–1534. DOI: 10.1007/s00425-006-0437-3

Romero G, Marino N, MacDonald A, et al. Srivastava D. 2020. Extreme rainfall events alter the trophic structure in bromeliad tanks across the Neotropics. Nature Communications 11: 3215. DOI: 10.1038/s41467-020-17036-4

Scharmann M, Grafe T. 2013. Reinstatement of Nepenthes hemsleyana (Nepenthaceae), an endemic pitcher plant from Borneo, with a discussion of associated Nepenthes taxa. Blumea 58: 8–12. DOI: 10.3767/000651913X668465

Scharmann M, Wistuba A, Widmer A. 2021. Introgression is widespread in the radiation of carnivorous *Nepenthes* pitcher plants. Molecular Phylogenetics and Evolution 163: 107214. DOI: 10.1016/j.ympev.2021.107214

Schöner C, Schöner M, Kerth G, Grafe T. 2013. Supply determines demand: influence of partner quality and quantity on the interactions between bats and pitcher plants. Oecologia 173: 191–202. DOI: 10.1007/s00442-013-2615-x

Schöner C, Schöner M, Kerth G, Suhaini S, Grafe T. 2015a. Low costs reinforce the mutualism between bats and pitcher plants. Zoologischer Anzeiger 258: 1–5. DOI: 10.1016/j.jcz.2015.06.002

Schöner M, Schöner C, Simon R, Puechmaille S, Ji L, Kerth G. 2015b. Bats are acoustically attracted to mutualistic carnivorous plants. Current Biology 14: 1911–1916. DOI: 10.1016/j.cub.2015.05.054

Thornham D, Smith J, Grafe T, Federle W. 2012. Setting the trap: cleaning behaviour of Camponotus schmitzi ants increases long-term capture efficiency of their pitcher plant host, Nepenthes bicalcarata. Functional Ecology 26: 11–19. DOI: 10.1111/j.1365-2435.2011.01937.x

