## Supplementary material for "Regulation of pitcher fluid volume and properties in six ecologically distinct Bornean *Nepenthes* species": fig. S1

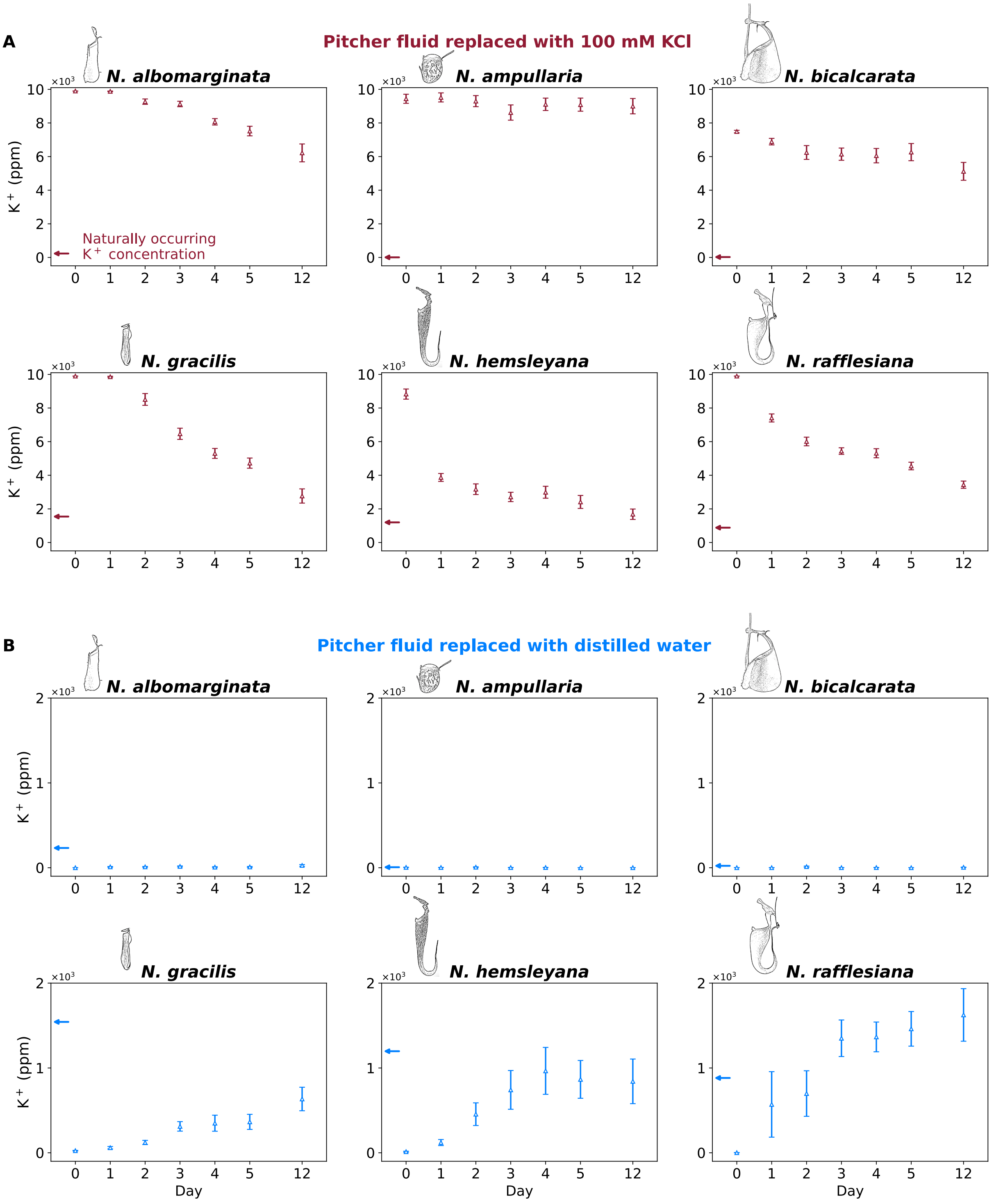


**Supplementary Figure 1:** Potassium concentration (open triangles) in pitcher fluids from six *Nepenthes* species following replacement with either (A) 100 mM KCl or (B) distilled water. Data are plotted as mean ± standard error (n=10 pitchers per species). Arrows indicate the mean naturally occurring potassium concentration for the particular species.
