## Supplementary material for "Regulation of pitcher fluid volume and properties in six ecologically distinct Bornean *Nepenthes* species": Table S1

<table>
<tr><th>Species</th><th>Pitcher height (H) (mm)</th><th>Fluid level (L) (mm)</th><th>Average relative pitcher fluid level (L/H) (%)</th><th>pH</th><th>Conductivity (mS/cm)</th><th>$K^+$ (ppm) (field measurement)</th><th>Surface tension (mN/m)</th><th>Extensional viscosity relaxation time (ms)</th></tr>
<tr><td>N. albomarginata</td><td>77.6 ± 17.6</td><td>27.2 ± 7.5</td><td>35.4 ± 7.9</td><td>4.6 ± 0.7</td><td>0.8 ± 0.3</td><td>382.6 ± 315.0</td><td>70.8 ± 0.8</td><td rowspan="3">No visible viscoelastic behaviour.</td></tr>
<tr><td>N. ampullaria</td><td>62.7 ± 8.3</td><td>28.2 ± 9.2</td><td>45.7 ± 15.1</td><td>5.9 ± 0.5</td><td>0.05 ± 0.02</td><td>0.0 ± 0.0</td><td>71.7 ± 1.0</td></tr>
<tr><td>N. bicalcarata</td><td>67.3 ± 11.1</td><td>44.9 ± 7.5</td><td>67.9 ± 13.3</td><td>5.3 ± 0.8</td><td>0.08 ± 0.02</td><td>84.1 ± 58.6</td><td>71.5 ± 0.5</td></tr>
<tr><td>N. gracilis</td><td>72.2 ± 13.8</td><td>17.2 ± 4.8</td><td>24.0 ± 6.1</td><td>3.1 ± 0.7</td><td>3.4 ± 1.0</td><td>860.4 ± 774.6</td><td>70.5 ± 1.3</td><td>0.45 ± 0.32</td></tr>
<tr><td>N. hemsleyana</td><td>126.6 ± 13.7</td><td>36.9 ± 15.6</td><td>28.7 ± 10.8</td><td>3.9 ± 0.9</td><td>1.6 ± 0.9</td><td>439.0 ± 338.4</td><td>69.0 ± 2.6</td><td>50.4 ± 29.8</td></tr>
<tr><td>N. rafflesiana</td><td>92.8 ± 16.3</td><td>57.1 ± 15.4</td><td>61.3 ± 10.3</td><td>2.9 ± 0.3</td><td>3.1 ± 1.3</td><td>1293.0 ± 709.7</td><td>65.7 ± 4.9</td><td>629.9 ± 200.5</td></tr>
</table>

**Table S1:** Field measurements of the level of pitcher fluid and of the fluid's physical and chemical properties in six *Nepenthes* species (same data as in Fig. 2). For each species, 10 fresh pitchers from separate plants were used for measurements and the mean values ± standard deviations are reported here.
