## Supplementary material for "Regulation of pitcher fluid volume and properties in six ecologically distinct Bornean *Nepenthes* species": Table S2

| | | Species → | *N. albomarginata* | *N. ampullaria* | *N. bicalcarata* | *N. gracilis* | *N. hemsleyana* | *N. rafflesiana* |
|---|---|---|---|---|---|---|---|---|
| **Treatment** | **Parameter** | **Comparison** | | | | | | |
| **100mM KCl** | **Fluid Level** | **Day 0 vs Day 1** | $t_9$=-0.8, p=0.43 | $t_9$=-1.8, p=0.11 | $t_9$=0.6, p=0.55 | $t_9$=-6.6, p<0.001 | $t_9$=-10.6, p<0.001 | $t_9$=-11.3, p<0.001 |
| | | **Day 0 vs Day 12** | $t_9$=-2.0, p=0.08 | $t_9$=-0.8, p=0.44 | $t_9$=-1.0, p=0.36 | $t_9$=-2.7, p=0.03 | $t_9$=-6.5, p<0.001 | $t_9$=-8.9, p<0.001 |
| | **Conductivity** | **Day 0 vs Day 12** | $t_9$=12.6, p<0.001 | $t_9$=8.0, p<0.001 | $t_9$=8.7, p<0.001 | $t_9$=47.2, p<0.001 | $t_9$=16.9, p<0.001 | $t_9$=35.1, p<0.001 |
| | | **Unmanipulated vs Day 12** | $t_9$=11.6, p<0.001 | $t_9$=15.1, p<0.001 | $t_9$=-3.4 p<0.01 | $t_9$=1.34, p=0.22 | $t_9$=3.3, p<0.01 | $t_9$=13.0, p<0.001 |
| | **$K^+$** | **Day 0 vs Day 12** | $t_9$=6.9, p<0.001 | $t_9$=0.7, p=0.47 | $t_9$=4.5, p<0.01 | $t_9$=16.9, p<0.001 | $t_9$=17.2, p<0.001 | $t_9$=30.1, p<0.001 |
| | | **Unmanipulated vs Day 12** | $t_9$=11.3, p<0.001 | $t_9$=19.7, p<0.001 | $t_9$=9.6, p<0.001 | $t_9$=2.4, p=0.05 | $t_9$=-2.4, p=0.05 | $t_9$=11.9, p<0.001 |
| | **Conductivity x Volume Product** | **Day 0 vs Day 12** | $t_9$=5.4, p<0.001 | $t_9$=5.8, p<0.001 | $t_9$=4.9, p<0.01 | $t_9$=3.5, p<0.01 | $t_9$=2.6, p=0.03 | $t_9$=4.5, p<0.01 |
| | | **Unmanipulated vs Day 12** | $t_9$=7.0, p<0.001 | $t_9$=9.2, p<0.001 | $t_9$=6.2, p<0.001 | $t_9$=2.5, p=0.04 | $t_9$=3.1, p=0.01 | $t_9$=6.9, p<0.001 |
| **Distilled Water** | **Fluid Level** | **Day 0 vs Day 1** | $t_9$=1.8, p=0.11 | $t_9$=-1.5, p=0.18 | $t_9$=1.1, p=0.30 | $t_9$=3.2, p=0.01 | $t_9$=6.5, p<0.001 | $t_9$=7.7, p<0.001 |
| | | **Day 0 vs Day 12** | $t_9$=4.6, p<0.01 | $t_9$=2.6, p=0.03 | $t_9$=1.6, p=0.14 | $t_9$=3.4, p<0.01 | $t_9$=0.3, p=0.77 | $t_9$=2.3, p=0.06 |
| | **Conductivity** | **Day 0 vs Day 12** | $t_9$=-2.6, p=0.04 | $t_9$=-3.8, p<0.01 | $t_9$=-3.7, p<0.01 | $t_9$=-10.6, p<0.001 | $t_9$=-6.8, p<0.001 | $t_9$=-12.3, p<0.001 |
| | | **Unmanipulated vs Day 12** | $t_9$=-12.4, p<0.001 | $t_9$=-0.3, p=0.80 | $t_9$=-3.4, p<0.01 | $t_9$=-2.2, p=0.06 | $t_9$=-0.3, p=0.80 | $t_9$=6.3, p<0.001 |
| | **$K^+$** | **Day 0 vs Day 12** | $t_9$=-2.6, =0.04 | $t_9$=1.5, p=0.18 | $t_9$=-1.5, p=0.18 | $t_9$=-4.4, p<0.01 | $t_9$=-3.1, p=0.01 | $t_9$=-5.2, p<0.001 |
| | | **Unmanipulated vs Day 12** | $t_9$=-19.8, p<0.001 | $t_9$=-14.1, p<0.001 | $t_9$=-7.0, p<0.001 | $t_9$=-6.6, p<0.001 | $t_9$=-1.4, p=0.21 | $t_9$=2.4, p=0.05 |
| | **Conductivity x Volume Product** | **Day 0 vs Day 12** | $t_9$=-1.9, p=0.09 | $t_9$=-1.1, p=0.29 | $t_9$=-2.3, p=0.06 | $t_9$=-2.9, p=0.02 | $t_9$=-8.9, p<0.001 | $t_9$=-8.2, p<0.001 |
| | | **Unmanipulated vs Day 12** | $t_9$=-54.9, p<0.001 | $t_9$=-9.7, p<0.001 | $t_9$=-9.5, p<0.001 | $t_9$=-6.9, p<0.001 | $t_9$=-4.6, p<0.01 | $t_9$=-5.4, p<0.001 |

**Table S2:** Statistical results of experiments testing the concentration dependence of fluid regulatory responses in six *Nepenthes* species (shown in fig. 5-7). In each case, paired t-tests are performed on the data listed in the 'Comparison' column.
