## Supplementary material for "Regulation of pitcher fluid volume and properties in six ecologically distinct Bornean *Nepenthes* species": Table S3

| Species | Day 0 vs Day 1 | | Unmanipulated vs Day 12 | |
|---|---|---|---|---|
| | $t_9$ | p | $t_9$ | p |
| *N. albomarginata* | 19.7 | <0.001*** | 0.9 | 0.388 |
| *N. ampullaria* | 18.7 | <0.001*** | -6.1 | <0.001*** |
| *N. bicalcarata* | 14.8 | <0.001*** | 0.2 | 0.884 |
| *N. gracilis* | 30.3 | <0.001*** | -0.2 | 0.838 |
| *N. hemsleyana* | 21.3 | <0.001*** | -3.9 | 0.004** |
| *N. rafflesiana* | 37.5 | <0.001*** | 1.0 | 0.338 |

**Table S3:** Paired t-test results for the pH of pitcher fluids measured after experimental alkalisation of the pitcher fluid with NaOH (bringing the pH to 11) and 24 hours after and measured before experimental manipulation versus 12 days after.
