## Supplementary material for "Regulation of pitcher fluid volume and properties in six ecologically distinct Bornean *Nepenthes* species": Table S4

| Pitcher response | Fluid absorption after filling with pitcher fluid | Fluid secretion after emptying | Fluid absorption after pitcher fluid replaced with distilled water | Fluid secretion after pitcher fluid replaced with 100mM KCl | Ion absorption after filling with pitcher fluid | Ion secretion after emptying | Ion secretion after pitcher fluid replaced with distilled water | Ion absorption after pitcher fluid replaced with 100mM KCl | pH change after alkalisation |
|---|---|---|---|---|---|---|---|---|---|
| **Fluid absorption after filling with pitcher fluid** | | r=0.24, p=0.644 | r=0.32, p=0.538 | r=0.19, p=0.713 | r=0.20, p=0.707 | r=0.12, p=0.817 | r=0.20, p=0.709 | r=0.50, p=0.308 | r=0.46, p=0.357 |
| **Fluid secretion after emptying** | | | r=0.90, p=0.015* | r=0.85, p=0.037* | r=0.24, p=0.652 | r=0.73, p=0.100 | r=0.34, p=0.516 | r=0.38, p=0.462 | r=0.70, p=0.126 |
| **Fluid absorption after pitcher fluid replaced with distilled water** | | | | r=0.84, p=0.037* | r=0.42, p=0.401 | r=0.89, p=0.017* | r=0.47, p=0.342 | r=0.38, p=0.464 | r=0.66, p=0.154 |
| **Fluid secretion after pitcher fluid replaced with 100mM KCl** | | | | | r=0.68, p=0.138 | r=0.91, p=0.012* | r=0.75, p=0.086 | r=0.53, p=0.277 | r=0.87, p=0.025* |
| **Ion absorption after filling with pitcher fluid** | | | | | | r=0.74, p=0.093 | r=0.99, p<0.001*** | r=0.32, p=0.542 | r=0.58, p=0.229 |
| **Ion secretion after emptying** | | | | | | | r=0.77, p=0.076 | r=0.38, p=0.458 | r=0.69, p=0.126 |
| **Ion secretion after pitcher fluid replaced with distilled water** | | | | | | | | r=0.32, p=0.534 | r=0.63, p=0.181 |
| **Ion absorption after pitcher fluid replaced with 100mM KCl** | | | | | | | | | r=0.87, p=0.025* |
| **pH change after alkalisation** | | | | | | | | | |

**Table S4:** Pearson correlation coefficients between rates of regulatory pitcher responses (calculated as the relative change during the first 24 hours) and their significance. Data used to generate species-wise means and standard deviations for correlations is the same as that used in figure 9.
